# Multi-Omic Analysis Reveals Disruption of Cholesterol Homeostasis by Cannabidiol in Human Cell Lines

**DOI:** 10.1101/2020.06.03.130864

**Authors:** Steven E. Guard, Douglas A. Chapnick, Zachary C. Poss, Christopher C. Ebmeier, Jeremy Jacobsen, Travis Nemkov, Kerri A. Ball, Kristofor J. Webb, Helen L. Simpson, Stephen Coleman, Eric Bunker, Adrian Ramirez, Julie A. Reisz, Robert Sievers, Michael H.B. Stowell, Angelo D’Alessandro, Xuedong Liu, William M. Old

## Abstract

The non-psychoactive cannabinoid, cannabidiol (CBD), is FDA-approved for treatment of two drug-resistant epileptic disorders, and is seeing increased use among the general public, yet the mechanisms that underlie its therapeutic effects and side-effect profiles remain unclear. Here, we report a systems-level analysis of CBD action in human cell lines using temporal multi-omic profiling. FRET-based biosensor screening revealed that CBD treatment resulted in a sharp rise in cytosolic calcium, and activation of AMPK and ERK kinases in human keratinocyte and neuroblastoma cell lines. CBD treatment led to alterations in the abundance of metabolites, mRNA transcripts, and proteins consistent with activation of cholesterol biosynthesis, transport and storage. We found that CBD rapidly incorporated into cellular membranes and altered cholesterol chemical activity, suggesting direct perturbation of cholesterol-dependent membrane properties. CBD treatment induced apoptosis in a dose-dependent manner in multiple human cell lines, which was rescued by inhibition of cholesterol synthesis, and potentiated by compounds that disrupt cholesterol trafficking and storage. Our data point to a pharmacological interaction of CBD with cholesterol homeostasis pathways, with potential implications in its therapeutic use.

## Introduction

The non-psychoactive cannabinoid, cannabidiol (CBD), was recently approved by the FDA for reducing convulsive seizure frequency in the drug-resistant epileptic disorders, Lennox-Gastaut syndrome and Dravet syndrome (Devinsky et al., 2017; Miller et al., 2020; Thiele et al., 2018). CBD lacks the psychotropic effects of its structural isomer, tetrahydrocannabinol, due to weak binding affinity (>3- 10 µM) for the endocannabinoid receptor CB_1_ (McPartland et al. 2015). CBD is well tolerated in humans (Iffland & Grotenhermen, 2017), and has been proposed as a potential therapeutic for a wide range of conditions, including colitis (De Filippis et al., 2011), cancer (Hinz & Ramer, 2019; Kenyon et al., 2018; Massi et al., 2013; McAllister et al., 2015), neuroinflammation (Esposito et al., 2011), chemotherapy associated nephrotoxicity (Pan et al., 2009), cardiomyopathy, and diabetic complications (Rajesh et al., 2010). Despite decades of research, the molecular mechanisms that underlie its therapeutic effects (Devinsky et al., 2014) and side-effect profiles (Devinsky et al., 2017, 2018; Thiele et al., 2018) are not well understood.

More than 65 protein targets of CBD have been proposed, 22 of which are membrane-localized channels and receptors (Ahrens et al., 2009; Ibeas Bih et al., 2015; Lauckner et al., 2008; Whyte et al., 2009). For example, CBD has been shown to inhibit voltage-dependent sodium currents mediated by the NaV1.1 sodium channel (Ghovanloo et al., 2018), mutations in which cause Dravet syndrome (Dravet, 2011). CBD has also been shown to inhibit voltage-dependent ion currents of six other human sodium channels, the Kv2.1 potassium channel, and even a bacterial sodium channel, with IC_50_ values 1-3 μM (Ghovanloo et al., 2018). Proposed targets also include calcium channels or receptors that regulate calcium, including T-type calcium channels (Ross et al., 2008), voltage-dependent anion channel 1 (VDAC1) (N Rimmerman et al., 2013), G protein-coupled receptor 55 (GPR55) (Lauckner et al., 2008), voltage-gated calcium channel Cav3.x (Ross et al., 2008), and transient receptor potential cation channels 1-4 (TRPV1-4) (Ibeas Bih et al., 2015). Postsynaptic calcium mobilization has been proposed as a mechanism to explain the anticonvulsant activity of CBD (Gray & Whalley, 2020). The ability of CBD to modulate many structurally diverse membrane channels and receptors raises the question of whether it acts through non-specific mechanisms, for example through biophysical alteration of lipid bilayers in which many of the proposed targets reside (Ghovanloo et al., 2018; Lundbaek et al., 2005; Lundbæk et al., 1996; Watkins, 2019).

Comparatively less is known about intracellular targets and pathways engaged by CBD in humans. In microglial cells, CBD has anti-inflammatory activity, and upregulates mRNA transcripts involved in fatty acid metabolism and cholesterol biosynthesis (Neta Rimmerman et al., 2011). In adipocytes, CBD leads to accumulation of triglyceride species, concomitant with phosphorylation changes of regulatory proteins controlling lipid metabolism, including CREB, AMPKA2 and HSP60 (Silvestri et al., 2015). In mice, CBD attenuates liver steatosis and metabolic dysregulation associated with chronic alcohol feeding (Y. Wang et al., 2017). These studies point to a systematic modulation of lipid and cholesterol pathways by CBD in animal models and human cell lines via as yet unknown mechanisms. The large-scale alteration in transcripts, proteins, and metabolites across numerous pathways suggests that CBD acts pleiotropically through numerous biomolecular targets and/or nonspecific effects on cellular membranes. This evidence motivated us to use an unbiased systems-based approach to examine the molecular basis of CBD cellular perturbation.

Recent advances in mass spectrometry has enabled comprehensive identification and quantification of cellular proteomes and metabolomes (Blum et al., 2018). Multi-omic profiling strategies that combine mass spectrometry based proteomics with transcriptome profiling can reveal critical and unexpected insights into the mechanisms of drug action (Hafner et al., 2019; Norris et al., 2017). In this study, we examined phenotypic and molecular responses to CBD treatment in a human neuroblastoma cell line, SK-N-BE(2), using 4 complementary experimental approaches: (1) high-content imaging of Förster resonance energy transfer (FRET) biosensors monitoring a panel of cellular activities, (2) subcellular proteomics, (3) phosphoproteomics, and (4) flux metabolomics. We found that CBD led to a chronic rise in cytosolic calcium and activation of 5’-AMP-activated protein kinase (AMPK) signaling within three hours post-treatment. In SK-N-BE(2) cells grown in cholesterol-replete media, CBD treatment led to increased abundance of mRNA transcripts and proteins involved in cholesterol import and biosynthesis. Metabolomics revealed a concomitant CBD-dependent increase in flux of glucose-derived carbon through cholesterol biosynthesis intermediates, despite being grown in cholesterol-replete conditions, suggesting that cholesterol sensing and synthesis become decoupled in the presence of CBD. We further show that CBD sensitizes human cells to apoptosis when co-treated with inhibitors of cholesterol trafficking and storage. Conversely, atorvastatin, an inhibitor of cholesterol biosynthesis, rescued cells from CBD-induced apoptosis. Together, our data reveal that CBD partitions into cellular membranes, and leads to disruption of cholesterol homeostasis and membrane-dependent processes.

## Results

### FRET-based sensor array reveals CBD response dynamics

To identify molecular events initiated by CBD treatment, we performed temporal multi-omic profiling of CBD treated human neuroblastoma cells. The dynamics of metabolite, RNA, and protein changes in response to drug perturbation could span time scales ranging from seconds to days, presenting a challenge for selecting appropriate time points in multi-omic analysis. To identify the optimal time points and CBD dose for multi-omic profiling, we used high-content imaging to monitor a panel of human cell lines (SK-N-BE(2) neuroblastoma and HaCaT keratinocyte cells) expressing FRET sensors. Transgenic lines were generated, each expressing a genetically encoded FRET biosensor gene capable of reporting a cellular activity (Chapnick et al., 2019). Sensors were selected to profile a broad range of activities, including abundance changes in metabolites and secondary messengers, as well as kinase and protease activities **(Table S1A).**

FRET ratios were measured in a time course following cells treated with vehicle or CBD across a range of doses from 0 to 100 µM **(Figure S1A)**. At each time point, we fit a log-logistic function with the FRET ratio data to estimate EC_50_ values and quantify the dose-dependency for each sensor over time. We found that cytosolic calcium, plasma membrane charge, AMPK kinase activity, ERK kinase activity, and glucose abundance exhibited the most significant dose-dependent changes. Several sensors showed time dependent responses at various doses of CBD but were eliminated for dose and time range analysis due to their poor dose-dependency (*R*^2^ <= 0.75). The EC_50_ distribution of CBD dose responses across all time points and biosensors displayed a median of 8.5 µM for SK-N-BE(2) cells. At early time points, a minimum of 20 µM CBD was required to activate the FRET sensors for which an EC_50_ could be estimated, including cytosolic calcium, AMPK activity, and plasma membrane charge **(Figure 1A).** SK-N-BE(2) cells displayed a higher degree of dose-dependency in FRET sensor activation over time relative to HaCaT cells **(Figure S1A).** We therefore selected SK-N-BE(2) cells treated with 20 µM CBD for subsequent multi-omic experiments.

**Figure 1.**
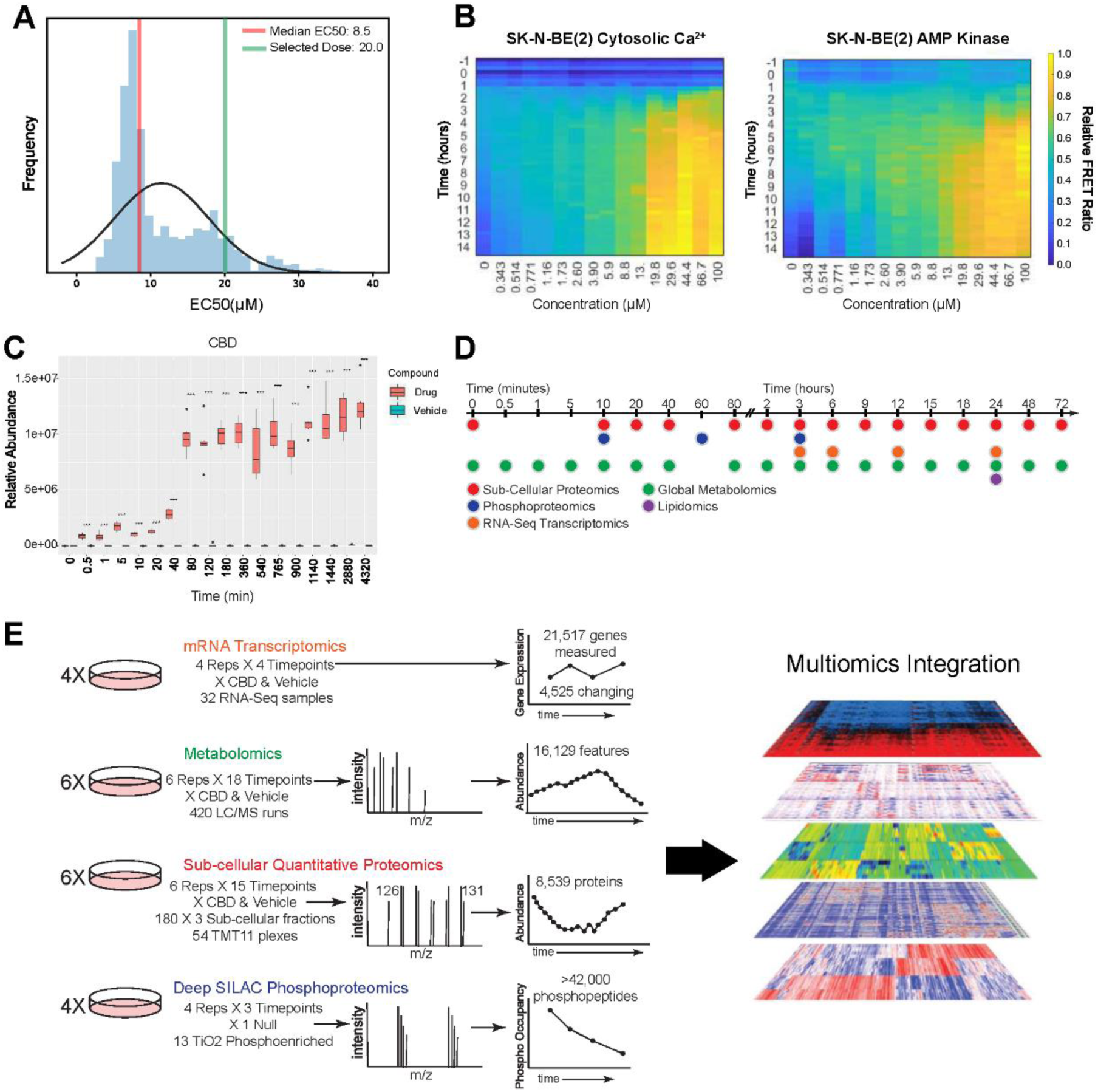
FRET biosensor screening strategy for dose and time selection of multi-omic CBD perturbation analysis. **(A)** EC_50_ distribution of CBD treatment across all sensors and time points (See also Table S1 for list of sensors). Dose response curves were fit to determine EC_50_ values (R^2^ values > 0.75) **(B)** Heat maps of FRET biosensor responses to CBD treatment for cytosolic Ca^2+^ and AMPK activity in SK-N-BE(2) cells, displaying FRET ratio over time at CBD doses from 343 nM to 100 µM. **(C)** Relative abundance of CBD over time from metabolomic profiling of SK-N-BE(2) cells treated with 20 µM CBD**. (D,E)** Time course schematic of multi-omic experimental strategy.

The biosensor data revealed that CBD led to activation of a diverse spectrum of cellular activities. After treatment with 20 µM CBD, the earliest dose-dependent events were increased cytosolic calcium at 3 hours followed closely by AMPK activation **(Figure 1B).** AMPK can be activated by distinct mechanisms: through allosteric binding of AMP, as a result of increased cellular abundance of AMP relative to ATP, or through phosphorylation-dependent activation by Ca^2+^/Calmodulin-dependent protein kinase kinase β (CaMKKβ) or STK11 (also known as LKB1) (Hawley et al., 2003, 2005; Shaw et al., 2004). CaMKKβ increases the activity of AMPK in a calcium-dependent manner through direct interactions with its kinase domain, driving downstream secondary calcium signaling events (Hawley et al., 2005; Woods et al., 2005). Our observation that CBD treatment leads to increased cytosolic calcium is consistent with previous reports of CBD-induced increase in cytosolic calcium, which was proposed to act through TRPM8, TRPV receptors, or Voltage-dependent T type receptors (Ibeas Bih et al., 2015; N Rimmerman et al., 2013).

We next monitored the kinetics of cellular uptake of CBD in SK-N-BE(2) cells. The relative abundance of intracellular CBD was quantified by mass spectrometry in a time course from 30 seconds to 72 hours. CBD was detected in cells as early as 30 seconds but did not reach steady state until 80 minutes post-treatment **(Figure 1C).** Based on these kinetics of CBD uptake, we performed a set of multi-omic experiments to examine the temporal response of SK-N-BE(2) cells to CBD treatment, from minutes to days, using global metabolomics, lipidomics, phosphoproteomics, subcellular proteomics, and transcriptomics **(Figure 1D),** resulting in the detection of > 42,000 phosphorylated peptides, 8,359 proteins, 21,517 gene transcripts and 16,129 metabolic features **(Figure 1E)**.

### CBD activates AMPK signaling and downstream substrate phosphorylation

We performed quantitative phosphoproteomics to quantify changes in phosphorylation in response to CBD treatment at 10 minutes, 1 hour, and 3 hours, using stable isotope labeling of amino acids in cell culture (SILAC) (Ong et al., 2002). At 10 minutes, only five significantly changing phosphorylation sites were observed (q < 0.05 and |log2 ratio| > 0.5) **(Figure S2A).** However, the number of significantly changing sites increased to 154 by 1 hour **(Figure 2A)**, mirroring the kinetics of CBD uptake into cells between 40 and 80 minutes **(Figure 1C)**. At both 1 hour and 3-hour time points, significantly changing phosphorylation sites were enriched in known effectors of AMPK signaling. **(Figure 2A, 2B and S2B).** The canonical phosphorylation motif of high confidence AMPK substrates has been identified as L-X-R-X-X-(pS/pT)-X-X-X-L (Dale et al., 1995; Gwinn et al., 2008; Schaffer et al., 2015). We found that AMPK motifs were significantly enriched in CBD-responsive phosphorylation sites at 1- and 3-hours, including L-X-R-X-X-pS and R-X-X-pS-X-X-X-L **(Figure 2C, 2D, and Figure S2C, S2D)**.

**Figure 2.**
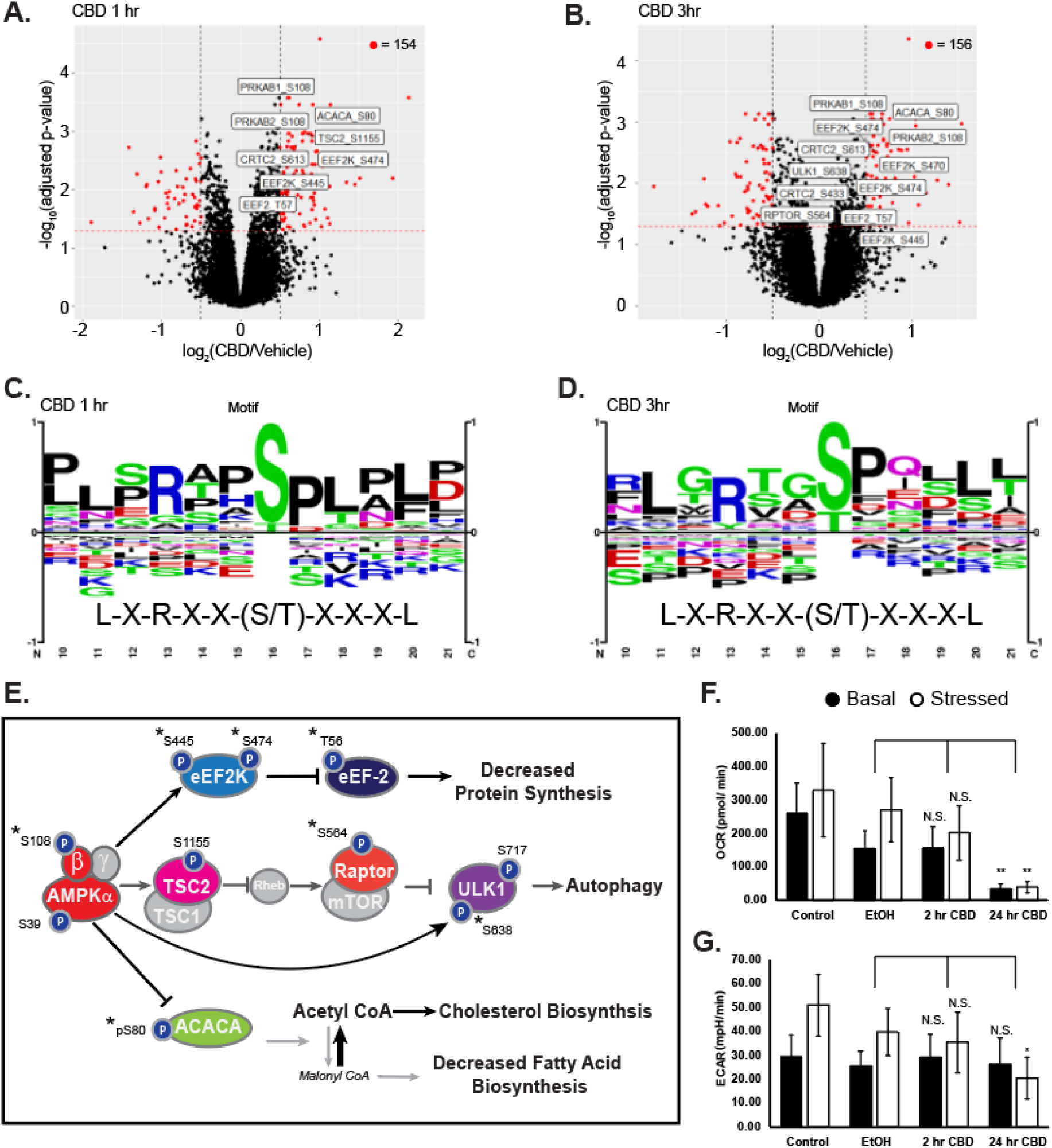
CBD increases phosphorylation of AMPK signaling proteins at early time points. (A-B) Volcano plots of quantified phosphorylation sites at 1 and 3 hours post treatment with 20 µM CBD, showing significance of differential change on y-axis and log_2_(CBD/DMSO) ratio on x-axis. Red: adjusted p < 0.05, |log_2_(CBD/DMSO) | > 0.5. Sites on AMPK proteins involved in AMPK signaling with adjusted p-value < 0.05 are annotated in white boxes. (C-D) Phospho-motif enrichment from phosphorylation sites identified as significantly changing in CBD treated cells vs vehicle control at 1- and 3-hours post CBD treatment. The AMPK averaged motif is displayed beneath the sequence logo (Schaffer et al., 2015) (E) Overlay of significantly changing phosphorylated sites onto an AMPK signaling diagram. Asterisks signify phosphorylation sites with a known upstream kinase (F-G) Seahorse extracellular flux measurement of oxygen consumption rate (OCR) and extracellular acidification rate (ECAR), * p < 0.05; ** p < 0.01. Stressed condition: Oligomycin treatment (1 µM)/ FCCP treatment (1 µM).

Of the CBD-responsive phosphorylation sites on proteins involved in AMPK signaling, several are annotated with biological function. We observed increased phosphorylation of S108 within the beta subunit of AMPK at the 1-hour and 3-hour time points. Phosphorylation of S108 drives a conformational change in the AMPK complex, resulting in stabilization of active kinase by preventing dephosphorylation of the activation site, T172 (Li et al., 2015). In agreement with increased AMPK activity after 1 hour of CBD treatment, we found increased phosphorylation of S80 on acetyl-CoA Carboxylase (ACACA), a known AMPK phosphorylation site **(Figure 2E)** (Carlson & Kim, 1973; Munday, 2002). ACACA catalyzes the rate-limiting step of fatty acid synthesis and is deactivated by AMPK phosphorylation of S80. Phosphorylation of ACACA S80 results in reduced conversion of acetyl-CoA into malonyl-CoA, reducing carbon flux through fatty acid synthesis, and increasing catabolic fatty acid β-oxidation (Fediuc et al., 2006; McFadden & Corl, 2009). In line with these findings, we observed decreased flux of carbon into *de novo* synthesized fatty acids **(Figure S2E)**. We found significantly decreased levels of short and medium chain, but not long chain acylcarnitines in the CBD-treated cells, indicating that fatty acid mobilization is comparable in the two groups, but more rapidly fluxed through fatty acid β-oxidation upon treatment with CBD **(Figure S2F)**.

We also identified increased phosphorylation of the translation elongation factor, EEF2, on T56 with CBD treatment at 1 and 3 hours. EEF2 T56 phosphorylation is sufficient to inhibit the GTP-dependent ribosomal translocation step during translational elongation, consistent with upstream activation of AMPK and EEF2K (Ryazanov et al., 1988). Together, these observations predict alterations of both protein and fatty acid synthesis downstream of AMPK activation by CBD.

Agreement between phosphoproteomics and the AMPKAR FRET biosensor data indicates that AMPK is activated by CBD treatment, raising the question of whether AMPK is activated through increased AMP:ATP ratio or by upstream kinases. To test whether CBD treatment acutely alters cellular energy status, we measured the oxygen consumption rate and extracellular acidification rate of CBD treated cells using a Seahorse extracellular flux assay. Treatment of SK-N-BE(2) cells with 20 µM CBD led to decreased levels of basal oxygen consumption by 24 hours, with little change at 2 hours post-drug treatment **(Figure 2F)**. Basal extracellular acidification rate remained unchanged **(Figure 2G)**. Consistently, we observed comparable rates of lactate production, but decreased carbon flux into TCA cycle metabolites in cells treated with CBD **(Figure S2G).** These results suggest that CBD-treated cells have decreased ATP production by mitochondrial respiration with little to no compensation by glycolysis, which may sustain AMPK activation at late time points. While we do not have direct evidence of the mechanism by which AMPK is activated between 1-3 hours, we hypothesize that in the absence of compromised ATP production at early time points, calcium influx into the cytoplasm may be responsible for activation of AMPK through upstream kinases such as CAMKKβ.

### CBD upregulates transcripts and proteins involved in cholesterol biosynthesis

To identify time-dependent proteome changes in subcellular compartments, we developed a pH-dependent cell fractionation scheme using differential centrifugation **(Figure 3A; Experimental Procedures).** The resulting ‘cytosolic’ fraction is enriched in soluble proteins from the cytosol, nucleus, and various luminal compartments (e.g. mitochondria) **(Figure S3A).** The first insoluble fraction, labeled as ‘membrane’, is enriched in proteins from mitochondrial and plasma membranes, while the second insoluble fraction is highly enriched in insoluble nuclear components, including condensed chromatin, spindles and nuclear speckles **(Figure S3B and S3C).** Principal component analysis (PCA) of these fractions revealed three compositionally distinct portions of the proteome, with each of these fractions exhibiting a time-dependent separation in response to CBD treatment. **(Figure S3D and S3E).** However, the membranous and nuclear fractions remain very similar in PCA space until 12 hour and later time points, suggesting relatively slow kinetics of protein regulation in response to CBD. Consistent with this observation, the frequency of significant events across fractions are limited at time points prior to 12 hours but increase dramatically to hundreds of proteins at time points between 15 to 72 hours. **(Figure 3B).**

**Figure 3.**
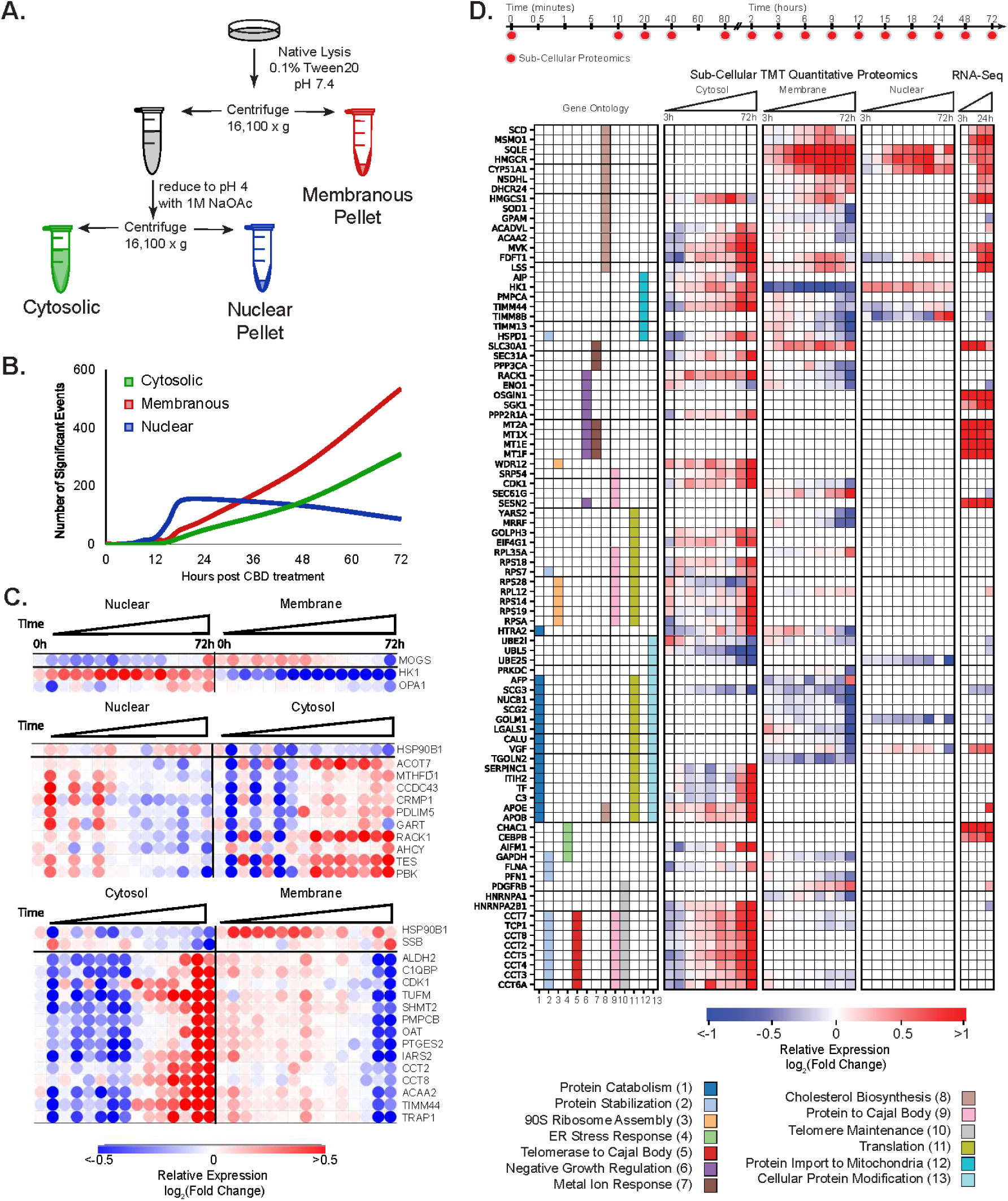
CBD treatment upregulates cholesterol biosynthesis enzymes and translocation of metabolic proteins. (A) Compositionally distinct subcellular proteomic fractions were fractionated by differential centrifugation and pH. The “cytosolic” fraction is enriched in soluble protein, the “nuclear” fraction is enriched in insoluble subnuclear compartments: condensed chromosome, spindles, spliceosomal complex etc., and the “membrane” fraction is enriched in membrane and mitochondrial related proteins. (See also Figure S3C -S3E) (B) Frequency of significantly changing proteomic events over time. (C) Anticorrelated proteins between proteomic fractions over time. PCA dimensionality reduction was used to decrease the impact of noisy signal contribution. Correlation between fractions, r < -0.8 was required. A large proportion of proteins listed above are known to compartmentalize in the mitochondria indicating protein shuttling or mitochondrial detachment/ attachment. (See also Figure S3F) (D) Proteins and mRNA transcripts that change significantly with CBD and map to the indicated gene ontology annotations that showed significant enrichment of differential proteins (Experimental Procedures).

A protein that translocated between cellular compartments in response to CBD would be expected have anticorrelated time courses in those subcellular fractions. To identify potential translocation events, we calculated the Pearson’s correlation coefficient between the temporal profiles of each of that protein’s subcellular fractions. We found 30 proteins with highly anti-correlated subcellular profiles **(Figure 3C).** Notably, Hexokinase1 (HK1) decreased in the membrane fraction and increased in the nuclear fraction **(Figure 3C and S3F).** HK1 detachment from the outer mitochondrial membrane attenuates conversion of the HK1 substrate glucose to glucose 6-P, decoupling glycolysis from mitochondrial respiration, and can alter the overall balance of energy metabolism in the cell (BeltrandelRio & Wilson, 1992; Crane & Sols, 1953; Saraiva et al., 2010). Consistently, CBD-treated cells exhibit reduced levels of glucose-6-phosphate at later time points **(Figure S3G).** This potential translocation event is consistent with decreased cellular respiration in response to CBD treatment **(Figure 2F)**, and previous reports of CBD-induced mitochondrial dysfunction in neuroblastoma cells (Alharris et al., 2019).

To identify CBD-dependent changes in mRNA transcript abundance, we performed an RNA-seq experiment, comparing SK-N-BE(2) cells treated with 20 µM CBD or vehicle for 3, 6, 12 and 24 hours. We identified 4,118 differentially expressed transcripts in CBD treated cells that were significant in at least one time point (q < 0.01). 204 of these genes displayed transcript abundances with a |log2 ratio| ≥1 **(Figure 1E)**. To identify potential transcription factor specific responses that explain mRNA transcript changes, we performed upstream regulator analysis on significantly changing transcripts (Krämer et al., 2014). The most enriched transcription factors for increasing transcripts shared oxidative stress as a stimulus and included ATF4, NFE2L2, SP1 **(Figure S3H)** (Blais et al., 2004; Dasari et al., 2006; Venugopal & Jaiswal, 1998; G. L. Wang & Semenza, 1993). CBD-treated cells also showed an accumulation of the principal cellular antioxidant glutathione, consistent with an oxidative stress response in CBD-treated cells **(Figure S3G).**

We merged differentially expressed transcript and protein identifications and performed gene ontology enrichment analysis using REVIGO pathway analysis **(Figure 3D)** (Supek et al., 2011). CBD responsive events were enriched in translation, endoplasmic reticulum (ER) stress response, metal ion response, and cholesterol biosynthesis (adj. p < 0.01). While many of these annotations are supported by either the transcriptome or proteome, dysregulation of cholesterol metabolism is supported by both. Within the cholesterol biosynthetic pathway ontology enrichment, 17 proteins displayed significant abundance changes that increased over time, including several key regulatory proteins. The rate limiting enzyme in cholesterol synthesis, HMGCR, increased by ∼300% on the protein level across both membranous fractions, with increased transcript abundance at 6 hours. Protein levels for SOD1, a negative regulator of HMGCR, decreased by 40%, consistent with derepression of *HMGCR* transcription (De Felice et al., 2004). The enzyme catalyzing the conversion of desmosterol into cholesterol in the terminal step in cholesterol biosynthesis, DHCR24, increased by 43% on the protein level in the “membrane” fraction (**Figure 3D**).

Together, the proteomic and transcriptomic data point to a concerted response in cholesterol homeostasis pathways and suggests that cells upregulate cholesterol biosynthesis capacity when challenged with CBD.

### CBD treatment results in accumulation of cholesterol biosynthesis intermediates and esterified cholesterol

Proteomic and transcriptomic analyses revealed a concerted CBD-induced upregulation of cholesterol biosynthesis machinery. These findings raised the question of whether CBD treatment leads to alterations in lipid and cholesterol metabolism (the latter pathway depicted in **Figure 4A**). We used mass spectrometry-based lipidomics to quantify the effect of CBD on lipids and sterols. Vehicle and CBD exposed cells were labeled with [U-^13^C_6_]-D-Glucose for 24 hours and harvested using methanol extraction. Cholesterol biosynthetic flux was quantified by mass spectrometry analysis of ^13^C incorporation into biosynthetic intermediates. We found that cholesterol precursors accumulated in CBD exposed cells **(Figure 4B and S4A)**, while labeled and total cholesterol itself decreased modestly (**Figure 4B and S4B**). This effect of CBD on total cellular cholesterol was confirmed using an Amplex Red cholesterol assay (**Figure S4C**).

**Figure 4.**
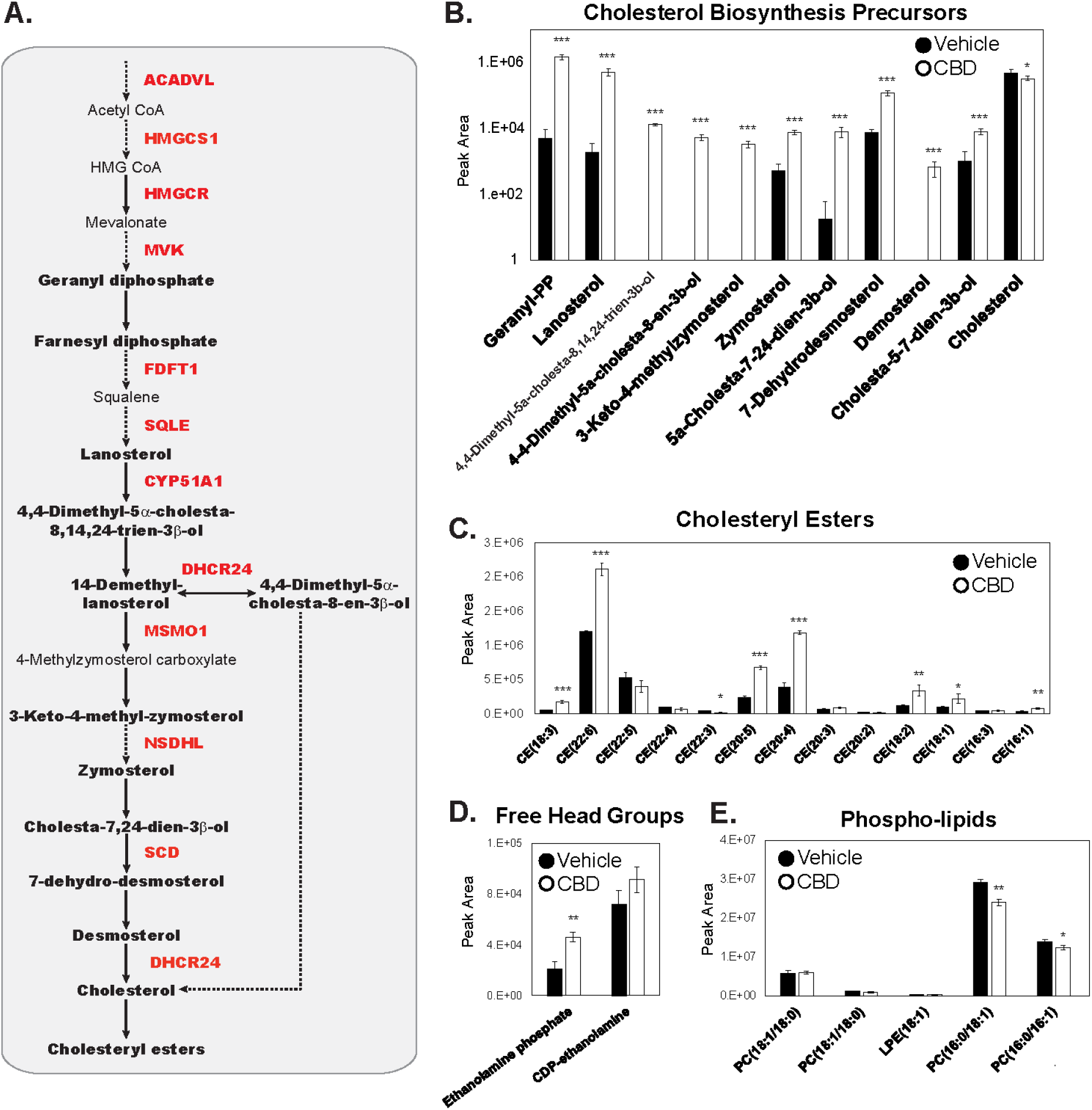
Cholesterol biosynthesis precursors and cholesterol esters accumulate upon CBD treatment. (A) Pathway diagram of cholesterol biosynthesis. Quantified metabolic intermediates are outlined in black bold, enzymes showing significant increase over time with CBD in proteomic analysis are shown bold red (Figure 3D). Dashed arrows indicate multiple intermediate steps that were not identified in the pathway. (B) D-glucose (U-^13^C_6_) metabolically labeled cholesterol biosynthesis precursors at 24 hours post 20 µM CBD treatment. (Student’s T Test : *p< 0.05, **p<0.01, ***p<0.001) (See also Figure S4) (C-E) Total abundance of lipids quantified by LC/MS/MS from lipid extracts of SK-N-BE(2) cells treated with vehicle or 20 µM CBD. Cholesteryl esters (CE), free head groups and phospholipids identified by mass spectrometry are displayed. Phosphatidylcholine(PC); Phosphatidylethanolamine (PE); lysophosphatidylethanolamine (LPE) (Student’s t-test: *p< 0.05, **p<0.01, ***p<0.001).

Internalized cholesterol is stored in lipid droplets after esterification with long-chain fatty acids by acyl-coenzyme A cholesterol O-acyltransferase enzymes (ACATs), using long chain fatty acyl-coenzyme A as the fatty acid donor (Brown et al., 1980). We detected accumulation of multiple species of cholesteryl esters with various chain lengths and acyl-chain saturation **(Figure 4C)**. Upregulation of cholesterol biosynthesis enzymes, together with increased abundance of metabolic precursors, suggest that CBD leads to increased production and storage of cholesterol esters. Due to the requirement of acyl-coenzyme A precursors in cholesterol esterification (T. Y. Chang et al., 2009), we surveyed our dataset for evidence of fatty acid utilization. Consistently, CBD treatment led to reduced levels of fatty acids after 48 hours (**Figure S4E).** We also found that a headgroup of sphingolipids, ethanolamine phosphate, increased by 2-fold in CBD treated cells, suggesting increased sphingolipid catabolism by the enzyme sphingosine-1-phosphate lyase (SGPL1) (**Figure 4D**) (Serra & Saba, 2010). We also surveyed the cellular abundance of all detectable species of phosphatidylcholine and phosphatidylethanolamine in cell extracts and found a variety of phospholipids that display reduced abundance in CBD treated versus vehicle treated cells (**Figure 4E**). Together, this evidence suggests that CBD leads to remodeling of the lipidome and perturbation to cholesterol homeostasis pathways.

### CBD increases storage and transport of cholesterol

Cholesterol is a critical structural component of cellular membranes. Alterations in cholesterol abundance can lead to severe cellular phenotypes that include mitochondrial dysfunction and apoptosis (Zhao et al., 2010). Disruption in cholesterol trafficking is a hallmark of Niemann-Pick disease type C (T.-Y. Chang et al., 2005), and can drive sodium channel-mediated inflammatory pain in animal models (Amsalem et al., 2018). To explore the implications of CBD’s disruption of cholesterol homeostasis, we tested whether CBD-induced cell death required upregulated cholesterol synthesis. Dose response analysis of cell viability revealed that 50% of SK-N-BE(2) cells die after 24 hours of treatment with 40 µM CBD (**Figure 5A**). SK-N-BE(2) cells were exposed to increasing concentrations of CBD in the presence or absence of the cholesterol biosynthesis inhibitor, atorvastatin, and analyzed for apoptosis using CellEvent caspase 3/7 dyes and live-cell fluorescence imaging. At 15 hours, 100 µM CBD leads to apoptosis of 50% of SK-N-BE(2) cells. Co-treatment of SK-N-BE(2) cells with CBD and atorvastatin reduced apoptosis by approximately 2-fold (**Figure 5B**). This atorvastatin-dependent rescue of CBD-induced apoptosis was far more pronounced in human HaCaT keratinocytes (**Figure S5A**), which are known to be highly sensitive to cholesterol perturbation (Bang et al., 2005). Further, CBD treated SK-N-BE(2) and HaCaT cells show an increase in apoptosis with increasing concentrations of a soluble form of cholesterol, 25-hydroxycholesterol (25-OHC) (**Figure 5C, and S5B).** Together, these results show that CBD sensitizes cells to apoptosis when challenged with excess cholesterol, either from endogenously synthesized or exogenous pools.

**Figure 5.**
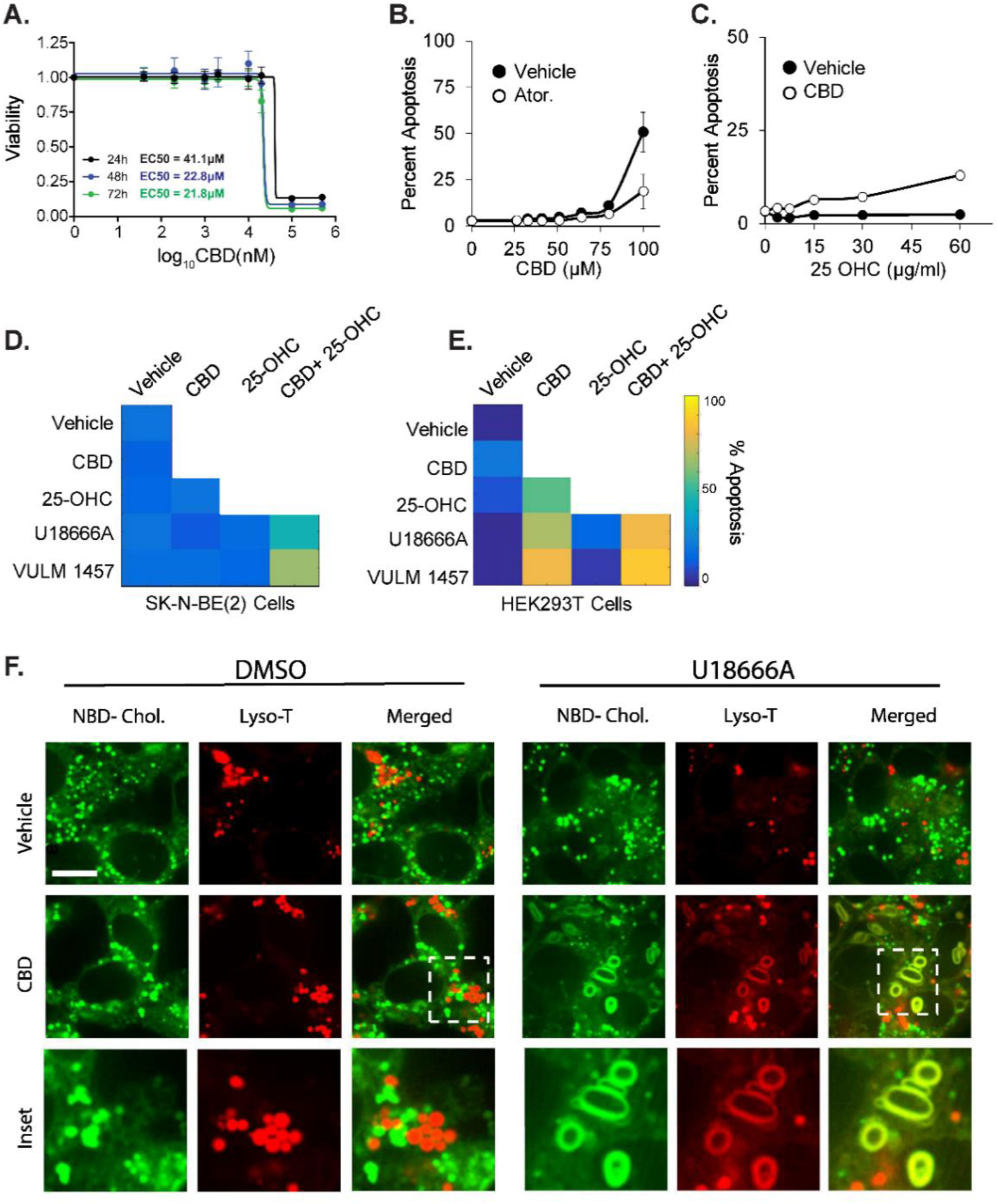
CBD induced apoptosis is rescued by inhibitors of cholesterol synthesis and increased by inhibitors of cholesterol transport and storage. (A) CBD was assessed for cytotoxicity in SK-N-BE(2) cells across increasing doses of CBD at 24, 48 and 72 hours by CellTiter-Glo luminescent assay. (B-E) SK-N-BE(2) or HEK293T cells were assessed for apoptosis at 24 hours using live cell microscopy using a resazurin based fluorometric cell viability stain. Cells were treated with 10 µM atorvastatin and exposed to increasing doses of CBD in B. 20 µM CBD and exposure to increasing doses of 25-OH cholesterol in C, and combinations of 20 µM CBD, 10 µM U18666A, 5µM VULM and 15µg/ ml 25-OH Cholesterol at 48 hours in D-E. Apoptosis displayed in a heatmap for each condition. (See also Figure S5) (F) Live cell confocal microscopy of SK-N-BE(2) with NBD-Cholesterol (Green) and a lysosomal dye Lyso-T (Red). Cholesterol subcellular distribution was examined upon exposure of cells to 20 µM CBD and/ or 10 µM U18666A. Scale bar: 3 µM

We next determined the dependency of CBD-induced apoptosis on cholesterol transport and storage. We measured apoptosis in SK-N-BE(2) and HEK293T cells treated with CBD and sublethal doses of 25-OHC (15 µg/ml), in combination with a cholesterol transport inhibitor (NPC1 inhibitor U18666A, 10 µM), or an inhibitor of ACAT, an enzyme required for esterification and intracellular storage of cholesterol (VULM 1457, 5 µM). Both compounds sensitized cells to apoptosis when CBD was present, which was more pronounced when cells were also challenged with 25-OHC (**Figure 5D and 5E**). VULM and U18666A treatment alone did not lead to increased apoptosis. These results demonstrate that interfering with cholesterol transport or cholesterol storage sensitizes cells to CBD-induced apoptosis.

One possible explanation for why CBD sensitizes cells to inhibitors of cholesterol trafficking and storage is that CBD increases the rate of cholesterol transport from the plasma membrane through the endosomal-lysosomal pathway. In support of this hypothesis, we observed increases in apolipoproteins B and E (APOB, APOE) abundance in the cytosol-enriched fraction. APOB/E are lipoprotein components of cholesterol-containing low-density lipoprotein particles required for cellular uptake of cholesterol **(Figure 3D)**. When cholesterol import through low density lipoprotein receptors (LDLR) is activated, and 25-OHC is supplied in excess, the inability to efficiently store cholesterol may cause accumulation of cholesterol in organelles that normally maintain low cholesterol levels. To examine this possibility, we visualized lysosomes (lysotracker dye) and cholesterol (NBD-cholesterol) in vehicle and CBD treated SK-N-BE(2) cells with live cell confocal microscopy.

In cells treated with either CBD or U18666A alone, puncta stained with lysotracker and NBD-cholesterol showed distinct spatial separation within cells. In contrast, co-treatment with CBD and U18666A led to formation of enlarged, membranous features co-stained with lysotracker and NBD-cholesterol (**Figure 5F**). Morphologically similar structures have been reported in models of Niemann-Pick disease type C, in which cholesterol accumulates in enlarged lamellar inclusions with components of lysosomes and endosomes, leading to a toxic cycle of enhanced cholesterol synthesis and intracellular accumulation (Demais et al., 2016; Höglinger et al., 2019). Sensitization of cells by CBD to compounds that disrupt intracellular trafficking of cholesterol, combined with our observations that CBD increases endogenous cholesterol biosynthesis and stimulates cholesterol esterification (**Figures 5D,E and 4B,C**), support a model where CBD increases transport of plasma membrane cholesterol through the endosomal-lysosomal pathway to intracellular compartments where it is esterified and sequestered, while escaping ER-resident cholesterol sensing machinery (Cheng et al., 1995; Y. Lange et al., 1999).

### CBD incorporates into membrane compartments and alters cholesterol accessibility and lateral diffusion

Subcellular fractionation of SK-N-BE(2) cells treated with CBD for 24 hours showed that CBD is concentrated primarily at the plasma membrane, with lower levels detected in ER and nuclear membranes (**Figure 6A**). CBD accumulation in the plasma membrane, and the CBD induced changes in cholesterol homeostasis suggest that CBD may alter cholesterol availability to sensing and trafficking proteins within the plasma membrane. To measure the effect of CBD on cholesterol accessibility, we measured the enzymatic oxidation rate of cholesterol to 5-cholesten-3-one by cholesterol oxidase in small unilamellar vesicles (SUVs). Cholesterol oxidase has been shown to sense alterations of lipid bilayer structure and cholesterol accessibility (Ahn & Sampson, 2004), and could reveal CBD-dependent alterations in cholesterol orientation in membranes. Titration of CBD into cholesterol-containing SUVs increased the initial reaction rate of cholesterol oxidase in a manner proportional to CBD concentration (**Figure 6B and S6A**).

**Figure 6.**
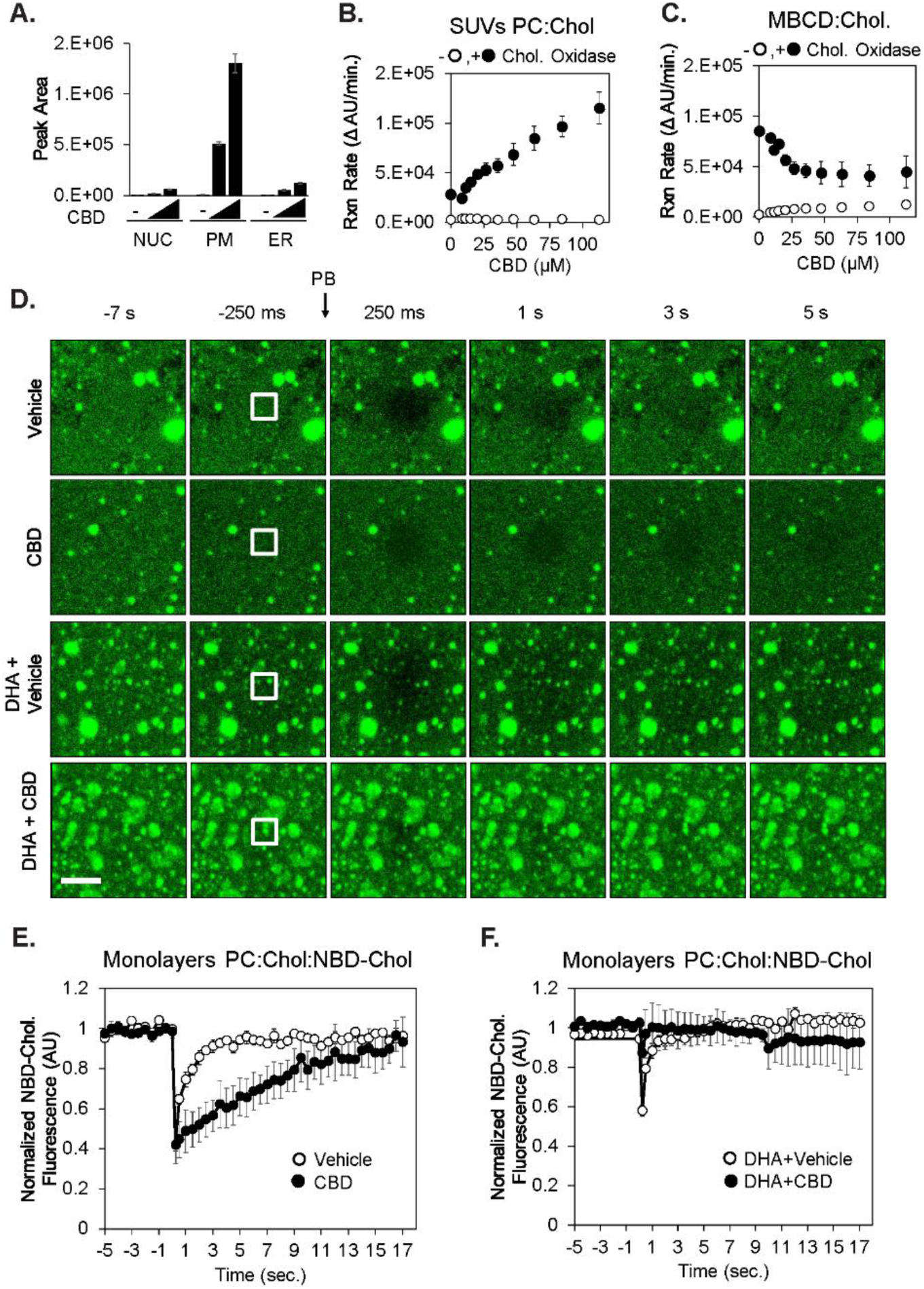
CBD incorporates into membranes, increases cholesterol chemical activity, and reduces lateral diffusion of cholesterol. (A) Ethanol extracts of subcellular fractions of SK-N-BE(2) cells exposed to 0, 20 and 40 µM CBD for 24 hr were analyzed for CBD using LC-MS. (B) Synthetic small unilamellar vesicles (SUVs-molar ratio: phosphatidylcholine:cholesterol:NBD cholesterol: 78:20:2 (n/n%) were used as a source of cholesterol in a fluorogenic cholesterol oxidase reaction to determine the effect of CBD on initial reaction rate. (C) Identical experiments were performed with cholesterol complexed to methyl beta cyclodextrin (MBCD), without SUVs present (D) Synthetic membrane monolayers containing NBD-cholesterol were adsorbed to borosilicate glass and used in fluorescent recovery after photobleaching (FRAP) experiments following exposure to either CBD (60 µM) and/or DHA (20 µM). Scale bar is 2.5 µm. Quantified fluorescence recovery after photobleaching is displayed in (E) and (F) (n=3).

The dose-dependent increase in cholesterol oxidase activity by CBD requires cholesterol localized in membranes. Freely soluble 25-OHC and soluble complexes of cholesterol and methyl beta cyclodextrin (MBCD) showed no dose dependence on CBD (**Figure 6C and S6B**). We repeated this experiment in a complex membrane environment using vesicles derived from ER membranes, and again observed a concentration dependent increase in cholesterol oxidase activity in response to CBD (**Figure S6C).** Together, these data provide evidence that CBD incorporates into membranes and alters cholesterol accessibility, likely by altering cholesterol orientation within the membrane to make the hydroxyl moiety more solvent accessible.

The ability of cholesterol oxidase assays to reveal alterations in lipid order has been previously reported in studies noting that cholesterol oxidase can preferentially target caveolar domains, a specialized type of lipid order domain (Ortegren et al., 2004; Smart et al., 1994). The ability of CBD to affect cholesterol activity in both synthetic and cell derived ER membranes suggests that CBD may contribute to changes in lipid order A hallmark of increased lipid order is a decrease in lateral diffusion of lipids (Ferreri, 2005; Lindblom & Orädd, 2009). We measured the effect of CBD on the lateral diffusion of fluorescently labelled cholesterol (NBD-cholesterol) in synthetic membrane monolayers. SUVs containing 20% (n/n%) of cholesterol and 2% (n/n%) NBD-cholesterol were deposited on glass-bottom multiwell imaging plates, followed by ultra-sonification. Recovery kinetics of fluorescent cholesterol were monitored in the presence of vehicle or CBD using fluorescence recovery after photobleaching (FRAP) **(Figure 6D)**. CBD significantly reduced the recovery of fluorescence in the photobleached monolayer area relative to vehicle control (**Figure 6D and 6E**), suggesting that CBD slows the lateral diffusion of fluorescent cholesterol. This effect of CBD on lateral diffusion could be rescued with simultaneous treatment of the docosahexaenoic acid (DHA), a known disrupter of lipid order (**Figures 6D and 6F**).

Our FRAP experiments demonstrate that DHA and CBD have opposing effects on the lateral diffusion of fluorescently labelled cholesterol in synthetic membranes However, it remains unclear how these biophysical effects of CBD and DHA on cholesterol impact cellular physiology. Esterification of DHA into membrane phospholipids results in remodeling of sphingolipid/ cholesterol-enriched lipid rafts, a known hub for apoptosis signaling (George & Wu, 2012; Wassall et al., 2018). To determine whether CBD and DHA also have opposing effects in a cellular context, we quantified the effect of CBD and DHA on cholesterol-dependent apoptosis. DHA treatment induced apoptosis in both HEK293T and SK-N-BE(2) cells in a dose dependent manner (**Figure S6D and S6E**), consistent with previous studies (Geng et al., 2018; Serini et al., 2008; Shin et al., 2013; Sun et al., 2013). Importantly, this DHA-induced apoptosis proved to be cholesterol dependent, as simultaneous treatment with DHA and the cholesterol sequestering agent, MBCD, delayed apoptosis in HEK293T cells, and fully rescued apoptosis in SK-N-BE(2) cells (**Figure S6D and S6E**). Similarly, CBD treatment (6.25 µM) rescued the apoptotic effects of DHA in both HEK293T and SK-N-BE(2) cells at 48 hours (**Figure S6D and S6E)**. These data indicate that CBD and DHA have opposing effects on cellular membrane structure and induction of apoptosis, both of which are cholesterol-dependent, but the connection between these two processes remain unclear.

Consistent with increased cholesterol accessibility, we found that CBD sensitized cells to permeabilization by the chemical agent filipin. Filipin is a highly fluorescent probe known to bind cholesterol and disrupt nearby lipid ordering (Schnitzer et al., 1994), resulting in permeabilization of membranes. Cells pretreated with 20 µM CBD for 24 hours were preferentially permeabilized by filipin, relative to vehicle control **(Figure S6F)**, further supporting a CBD-dependent alteration in plasma membrane structure and cholesterol. This data suggests that CBD either directly increases cholesterol availability to filipin, or destabilizes the membrane, thereby contributing to the membrane disruption effects of filipin.

## Discussion

Although clinical and preclinical evidence point to CBD as a promising therapeutic compound for epilepsy, the cellular targets that mediate its effects in humans remain unclear. In this study, we found that CBD elicited pleiotropic effects on the proteome, transcriptome and metabolome of human cells. Our data suggest that CBD integrates into cellular membranes and alters cholesterol orientation within the phospholipid environment. Partitioning of CBD into model membranes decreased lateral diffusion of cholesterol, altered cholesterol chemical activity, predicting that CBD may alter the biophysical properties of cellular membranes, with consequent effects on diverse membrane proteins and their downstream targets.

We found that CBD treatment led to increased cytosolic calcium within 2 hours in human neuroblastoma and keratinocyte cells. AMPK activity followed the observed increase in calcium, suggesting upstream activation of AMPK by the calcium-dependent kinase CAMKKβ. Compromised ATP generation by mitochondrial respiration may sustain AMPK activation after 24 hours, as suggested by our Seahorse analysis. Increased AMPK activity, and increased phosphorylation of its substrate ACACA, predicted reduced fatty acid synthesis and altered Acetyl-CoA metabolism, which was confirmed by metabolomics analysis. Upregulation of cholesterol biosynthesis on proteome, transcriptome and metabolomic levels occurred as early as 3 hours and was sustained up to 72 hours. As acetyl-CoA is precursor of cholesterol, the increase in cholesterol biosynthesis precursors are consistent with acetyl-CoA supporting this flux. Parallel to upregulation of cholesterol biosynthesis, we hypothesize that increased cholesterol import may occur through the LDLR-endocytic pathway, resulting in increased transport of cholesterol through endosomal-lysosomal trafficking. Increased stress on cholesterol trafficking and regulatory processes, combined with compromised cellular energetics driven by CBD may contribute to increased apoptosis in CBD-treated cells.

### Concordance of multi-omic data points to CBD disruption of cholesterol homeostasis

Integration of our transcriptomics, metabolomics, and proteomics data provided multiple lines of evidence for the disruption of cellular cholesterol homeostasis by CBD. Multiple aspects of cholesterol regulation were dysregulated by CBD: cholesterol biosynthesis (**Figure 3D and 4B**), transport (**Figure 3D and 5F**), and storage (**Figure 4C**). All three omics analyses provided evidence for perturbed cholesterol biosynthesis. For instance, transcriptomics and proteomics reported transcriptional activation and protein accumulation of the rate limiting enzyme in the biosynthetic pathway of cholesterol, HMGCR (**Figure 3D**). Increased HMGCR protein production is a canonical response to decreased cholesterol levels in the ER, where cholesterol is sensed through the SREBP-SCAP axis (Brown & Goldstein, 1997). Consistent with this observation, we found that cholesterol precursors accumulated in CBD-treated cells (**Figure 3B, S3A**), with a modest reduction in total cholesterol (**Figure 3B, S3A, S3B**) and a large increase in cholesterol esters.

The upregulation of cholesterol biosynthesis by CBD is paradoxical; cells cultured in cholesterol-replete media with abundant intracellular stores of esterified cholesterol typically downregulate cholesterol biosynthesis by the ER-resident SREBP sensing machinery to maintain homeostatic levels of cellular cholesterol (Brown & Goldstein, 1997). These results suggest that in the presence of CBD, the ER is unable to accurately sense the abundance of cholesterol at the plasma membrane, and as a result, generates an excess of cholesterol that is esterified. One possibility is that CBD prevents cholesterol sensing by GRAM domain proteins, which localize to plasma membrane- ER contact sites, bind and transport specific lipids between the two membranes (Besprozvannaya et al., 2018). Within this family, GRAMD1s sense and bind the ‘accessible’ pool of cholesterol that is not currently complexed with other lipid species, and transports it to the ER (Naito et al., 2019). The pools of cholesterol that are either ‘accessible’ or ‘inaccessible/sequestered’ are regulated by the domains they associate with, and are frequently driven by sphingomyelin and phospholipid association (Das et al., 2014; Yvonne Lange et al., 2013; Sokolov & Radhakrishnan, 2010). CBD driven alterations of cholesterol orientation and decreased lateral diffusion presented here, together with previously reported CBD-dependent increases in lipid raft stability and size, suggest that the pool of ‘sequestered cholesterol’ is increased in CBD treated conditions. However, specific investigation of GRAMD1 sensing and partitioning of CBD and cholesterol within the ER membrane will need to be investigated in future studies.

### The effects on cholesterol homeostasis in therapeutic applications of CBD

Biosensor and multi-omic profiling revealed the activation of a diverse spectrum of cellular activities by CBD; in particular, the disruption of cholesterol and lipid homeostasis that are important for proper membrane function. These results may have broad implications on the mechanistic underpinnings of the clinical effects of CBD. Transmembrane proteins known to be regulated through lipid ordered domains have been implicated in many of the diseases for which CBD has been proposed as a therapeutic. These include inflammatory disorders, Alzheimer’s disease, and several types of cancer (Gianfrancesco et al., 2018; Hsu et al., 2018; Mollinedo & Gajate, 2015; Pirmoradi et al., 2019; Staneva et al., 2018). Many targets of CBD proposed to underlie its efficacy as an anti-convulsant are also membrane proteins, including TRPV1, GPR55, and adenosine transport proteins (Bisogno et al., 2001; Liou et al., 2008; Ryberg et al., 2007). CBD inhibits ion currents from many structurally diverse voltage-gated ion channels at similar micromolar concentrations, and with a high degree of cooperativity, suggesting that CBD acts indirectly on ion channels through perturbation of membrane structure (Ghovanloo et al., 2018). Moving forward, it will be important to determine the role of lipid order and cholesterol orientation in the mediation of CBD-induced effects in models of generalized seizures.

Our study suggests that not all CBD effects on cells are therapeutically beneficial, and that high dose use of CBD may lead to cholesterol-dependent side effects in certain cell types that rely on high levels of cholesterol synthesis or import. We demonstrated that CBD driven apoptosis is heavily dependent on the cholesterol status of cells (**Figure 5B, 5C, S5A and S5B**). As a large fraction of cellular cholesterol in humans is synthesized in hepatocytes, we predict that many of the side effects of heavy CBD consumption may occur in the liver. Adverse events in CBD clinical trials include elevated liver aminotransferase levels (Gaston et al., 2017), a hallmark of liver injury (Giannini et al., 2005), which suggests that CBD may elevate the risk of hepatoxicity. Our data point to the importance of testing whether CBD use may interact adversely with certain dietary behaviors that elevate blood cholesterol, as the combination of cholesterol/ hydroxycholesterol and CBD is toxic to a cell line derived from skin (HaCaT cells), brain (SK-N-BE(2) cells), and kidney (HEK293T cells) (**Figures 5D, 5E, S5B**). Further, our results that CBD disrupted cholesterol trafficking through lysosomes, when in combination with U18666A, raises the question of whether CBD use might increase the risk of toxicity in patients with Niemann-Pick disease type C, which harbor mutations in NPC1, the target of U18666A (Lu et al., 2015).

## Experimental Procedures

### Compound Preparation

CBD was derived from domestically grown industrial hemp that was cultivated and purified by Sievers Infinity, LLC, Colorado-owned corporation, registered with the Colorado Department of Agriculture (CDA) to grow and cultivate industrial hemp (CDA # 69096). The purified hemp-derived material was characterized by mass spectrometry, X-ray diffraction, differential scanning calorimetry, nuclear magnetic resonance (1H-NMR and 13C-) spectroscopy, and HPLC-UV. The quantitative proton NMR results indicate that the sample is >95% CBD and the HPLC results indicate that 12 other commonly found cannabinoids (including THC) were less than the limit of detection of 0.004%.

### Generation of FRET Biosensor cell lines

Stable transgenic biosensor-expressing cell lines were made in HaCaT and SK-N-BE(2) cells as previously described (Chapnick et al., 2015). Briefly, biosensor gene-containing plasmids were obtained through the Addgene plasmid depository, and subcloned into our Bsr2 parent plasmid (sequence available upon request). Each biosensor Bsr2 plasmid was co-transfected with a PB recombinase expressing vector (mPB) via polymer-based transfection using polyethyleneimine (PEI) (Polysciences, 25kD Linear). Each stable transgenic cell line was selected for 7 days using 10 µg/ml Blasticidin S. FRET biosensor profiling was conducted in multiplexed parallel live cell experiments using 384 well imaging plates (Corning #3985) in an ImageXpress MicroXL high throughput microscope. Filters used for FRET measurements were the following: FRET excitation 438/24-25, dichroic 520LP, emission 542/27-25 (Semrock MOLE-0189); CFP excitation 438/24-25, dichroic 458LP, emission 483/32-25 (Semrock CFP-2432B-NTE-Zero). Time lapse microscopy images were collected, and FRET Ratio calculations for each site in each well at each time were performed as the mean value from pixels above threshold of background and flatfield corrected images, where each pixel value represented the FRET channel intensity divided by the CFP channel intensity. This method is described in more detail in our previous studies (Chapnick et al., 2015; Chapnick & Liu, 2014). Calculation and data visualization were performed in MATLAB using custom scripts that are available upon request.

### EC50 estimation from FRET sensor dose responses

Dose responses at each time point were fit with the following fit function y = 1 / (1 + np.exp(-k*(x-EC_50_))) using python’s scipy.optimize.curve_fit package. Prior to fitting, measurements were scaled between zero and one. R^2 goodness of fit was calculated between the sigmoid fit and the median of the replicates (duplicates) for each sensor/ time point combination. Fits with EC_50_ estimates outside the dose range were discarded. EC_50_ values were kept for fits that resulted in a R^2 GOF > 0.75. The resulting distribution of EC_50_ values was somewhat bimodal resulting in a median EC_50_ of 8.48 μM across all sensors and times.

### Transcriptomics Workflow

CBD, and vehicle treatments were prepared in quadruplicate (4 drug treated/ 4 vehicle controls) at 3, 6, 12, and 24-hour time points. 500 ng of Total RNA was used in Illumina TruSEQ mRNA library prep protocol. Libraries were run on the Illumina HiSEQ4000 at single read 150 bp length. Sequencing was performed on seven consecutive lanes. Median read counts per lane were ∼49,000 with a CV of ∼7%. Starting with 228 fastq files, each lane set was concatenated per condition. Run specifications were, 51 bp reads, standard single-read, first-stranded. Alignment to the human genome (HG19), was done using Tophat version 2.0.13. Two mismatches were allowed, and duplicate mapping was permitted at up to two locations. Using Trimmomatic-0.36, base pairs were trimmed if they dropped below a PHRED score of 15 within a sliding window of 4bp. Remaining reads shorter than 35 base pairs were removed. Illumina adapter sequences were also clipped using Trimmomatic. Fastqc was used to verify data integrity before and after trimming. Cufflinks/Cuffdiff 2.2.1 was used to obtain FPKM normalized gene count and differential expression measurements at each time point (Trapnell et al., 2010). One p/q-value was generated for each gene at each time point. Genes with a q-value significance < 0.05, and absolute log_2_ fold change of 1 or greater, for at least one time point, were retained for downstream analysis.

### Proteomics

#### Experimental Design and Statistical Rationale

Protein quantification for time-series was performed with a Tandem Mass Tag (TMT) isobarically labeled 11-plex multiplexing scheme. The 15-point time series for each cellular fraction was split into three series, with every series containing 5 treatment and matched control time point pairs, with 0 sec, 40 min, 3 hr, 12 hr, and 24 hr time points in Series A; 10 min, 80 min, 6 hr, 15 hr, and 48 hr in Series B; and 20 min, 2 hr, 9 hr, 18 hr, and 72 hr time points in Series C. This separation was performed so that a protein could be missing from one and/or two series due to stochastic effects in data-dependent acquisition and the overall trend could still be inferred, though with reduced resolution. The 11th label in each series was devoted to a global mix reference channel, which would be present in all series for a given cellular fraction. The global mix is a cell-fraction specific mixture that contains an equal portion from each time point sample. This channel was the denominator in the intermediate ratiometric measurement for differential expression for both drug-treated samples and time-matched controls. This mixture channel was constructed so that every measurable protein observed at any time point has a non-zero denominator when ratios are taken. When the differential expression is compared between the drug-treated labeled samples and matched control samples and expressed as a log_2_ ratio, the global mix reference channel cancels out.

The differential expression of each individual protein was determined using Bayesian methods for isobaric labeled proteomics data (Jow et al., 2014). Briefly, all observed peptides are mapped to a list of observed protein ID’s via Isoform Resolver (Meyer-Arendt et al., 2011). The TMT 11-plex reporter ion spectrum peaks for each peptide contributes to the inference of the differential expression of a protein and reporter ion label. In this case, each reporter ion label represents a measured time point. The label-to-label normalization is handled via a hierarchical model, which calculates the bias inherent with each specific label by pooling differential expression estimates from all proteins, changing and unchanging. The hierarchical models are solved computationally with a Markov Chain Monte Carlo (MCMC) method, running chains in parallel for faster results (Denwood, 2016). The MCMC returns a Gaussian posterior probability distribution of log_2_ differential expression for each protein for each label. The model initially fits the ratiometric differential expression for every treatment and matched control relative to a global mix channel, and the reported drug-induced differential expression is the difference (in log_2_ space) between the treated sample and the matched control sample. Five MCMC chains were run independently for at least 500k steps, and convergences for each run were verified via Gelman-Rubin convergence variable < 1.05 (Gelman & Rubin, 1992).

The differential expression was calculated independently for all biological replicates so protein-level variance from separate replicates could be examined and quantified in the posterior distributions obtained from MCMC. For reporting a single differential expression for a protein and label, the Bayesian updating procedure is used to produce a single posterior distribution, from which a mean point estimate and 95% credible interval are calculated. In some specific instances, labels represent technical rather than biological replicates. In cases of technical replicates, the point estimate values were averaged, and the credible intervals extents were treated as errors and added in quadrature. With this procedure, technical replicates contribute a single probability distribution to any further Bayesian updating.

For every cellular fraction and time point, then, there are between 3 and 6 biological replicates, and the number of replicates represented in the drug treated samples and the matched control samples are not necessarily the same. The effect size (Cohen’s *d*) was calculated between the posterior probability distributions of the drug treated and matched control samples as a standardized measure to determine if there was a drug effect. Statistical power analysis was performed to show that, with significance criteria α = 0.05 and the statistical power requirement (1-β) = 0.8, the appropriate effect size threshold should be *d* > (1.50, 1.75, 2.25, 3.38) for proteins observed within 6, 5, 4, or 3 replicates, respectively. A protein was selected for further consideration if it showed differential expression greater than this threshold for any given time point.

The Bioconductor Edge package (Leek et al., 2006) (DOI:10.18129/B9.bioc.edge) version 2.8.0 was used for time course differential analysis. Many proteins were not present for all replicates and/or plexes, so Edge was run sequentially to generate *p*-values for each case. For instance, in the soluble fraction, there were 273 proteins that were only present in two replicates. These were run through Edge separately from the other 1957 proteins that were observed in three replicates. The resulting time series *p*-values were combined into a list and FDR corrected using Benjamini-Hochberg multiple hypothesis correction (Benjamini & Hochberg, 1995).

### Proteomics Network Analysis

Of the significantly changing proteins, correlation networks were generated for each subcellular fraction. Networks were created from the ethanol (vehicle) treated samples, as well as for the CBD treated samples. Network edge values were assigned using Spearman correlation coefficients between all proteins (vertices) for a given replicate. For each pair of proteins, 2*N edge values were generated, where N is the number of available replicate measurements for that protein. An independent *t*-test was used between basal replicate edge values and treatment edge values to evaluate what edges were significantly changed due to CBD treatment. Edges with -log_10_(*p*-value) > 2 (*p* < 1%) were retained. Python graph-tool package was used to generate a stochastic block-model representation of the resulting network, which clusters nodes based on network connectivity similarity.

Combined Heatmap Criteria: All protein IDs with edge adjusted p-values less than 1% were merged with gene IDs from RNA-seq with edge adjusted p-value < 1% and minimum absolute log_2_ fold change > 0.5. Merged list was used as input for Enrichr (Kuleshov et al., 2016) to get a table of GO terms (go_biological_processes_2017). GO terms were reduced using REVIGO with “medium” size setting: Terms with dispensability score less than 0.1 and q-value < 5% were kept. Merged IDs from remaining GO ontologies were clustered and plotted in heatmap by relative expression in CBD treated condition compared to vehicle control at each time point starting at 3 hours.

### Subcellular Fractionation in Proteomics

For each sample, a 10 cm Petri Dish containing 10^6^ SK-N-BE(2) cells was harvested and washed three times with 10 ml of 20°C PBS. All PBS was removed by aspiration and plates were frozen using liquid nitrogen and stored at -80°C overnight. Each plate was thawed on ice and 400 µl Tween20 Buffer (1x PBS, 0.1 % Tween20, 5 mM EDTA, 30 mM NaF, 1 mM NaVo4, 100 µM Leupeptin, 2.5 µM Pepstatin A) and scraped thoroughly using a standard cell scraper. The resulting lysate was homogenized with a 200 µl pipette and transferred to 1.7 mL Eppendorf tube on ice. Lysate tubes were incubated for 30 min at 4°C rotating end-over-end. After rotation, tubes were centrifuged for 10 min at 4°C (16,100 rcf). All supernatant was transferred into new labeled 1.7 mL Eppendorf. This tube contains insoluble buoyant plasma membrane and cytosol. The leftover pellet is the ‘Membrane’ fraction and is enriched in nuclei. 40 µL of 1 M NaOAc was added to the supernatants, which immediately were exposed to centrifugation for 10 min at 4°C (16,100 rcf). All supernatant was transferred into new labeled 1.7 mL Eppendorf. This is the ‘Soluble’ fraction. The pellet was resuspended in 400 µl 20 °C SDS buffer. This is ‘Insoluble #2’ fraction. All fraction containing tubes were filled completely with -20 °C Acetone and stored overnight in -20 °C. Each tube was exposed to centrifugation for 10 min at 4°C (16,100 rcf) and supernatants were aspirated and discarded, while pellets were allowed pellets to air dry 10 min 20 °C. The pellets then proceeded to the FASP procedure.

### Quantitative Subcellular Proteomics

#### Sample preparation

Precipitated and dried subcellular protein extracts were solubilized with 4% (w/v) sodium dodecyl sulfate (SDS), 10 mM Tris(2-carboxyethyl)phosphine (TCEP), 40 mM chloroacetamide with 100 mM Tris base pH 8.5. SDS lysates were boiled at 95°C for 10 minutes and then 10 cycles in a Bioruptor Pico (Diagenode) of 30 seconds on and 30 second off per cycle, or until protein pellets were completely dissolved. Samples were then cleared at 21,130 x g for 10 minutes at 20⁰C, then digested into tryptic peptides using the filter-aided sample preparation (FASP) method (Wiśniewski, 2016). Briefly, SDS lysate samples were diluted 10-fold with 8 M Urea, 0.1 M Tris pH 8.5 and loaded onto an Amicon Ultra 0.5 mL 30 kD NMWL cutoff (Millipore) ultrafiltration device. Samples were washed in the filters three time with 8 M Urea, 0.1 M Tris pH 8.5, and again three times with 0.1 M Tris pH 8.5. Endoproteinase Lys-C (Wako) was added and incubated 2 hours rocking at room temperature, followed by trypsin (Pierce) which was incubated overnight rocking at room temperature. Tryptic peptides were eluted via centrifugation for 10 minutes at 10,000 x g and desalted using an Oasis HLB cartridge (Waters) according to the manufacture instructions.

### High pH C18 fractionation of TMT labeled peptides

Dried 10-plexed samples were then suspended in 20 uL 3% (v/v) acetonitrile (ACN) and 0.1% (v/v) trifluoroacetic acid (TFA) and loaded onto a custom fabricated reverse phase C18 column (0.5 x 200 mm C18, 1.8 μm 120Å Uchrom (nanoLCMS Solutions) maintained at 25⁰C and running 15 μL/min with buffer A, 10 mM ammonium formate, pH 10 and buffer B, 10 mM ammonium formate, pH 10 in 80% (v/v) ACN with a Waters M-class UPLC (Waters). Peptides were separated by gradient elution from 3% B to 50% B in 25 minutes, then from 50% B to 100% B in 5 minutes. Fractions were collected in seven rounds of concatenation for 30 s per fraction, then combined for a final of six high pH C18 fractions. Samples were dried and stored at -80⁰C until ready for LC/MS analyses.

### Liquid Chromatography/Mass spectrometry analysis

Samples were suspended in 3% (v/v) acetonitrile, 0.1% (v/v) trifluoroacetic acid and direct injected onto a 1.7 μm, 130Å C18, 75 μm X 250 mm M-class column (Waters), with a Waters M-class UPLC or a nanoLC1000 (Thermo Scientific). Tryptic peptides were gradient-eluted at 300 nL/minute, from 3% acetonitrile to 20% acetonitrile in 100 minutes into an Orbitrap Fusion mass spectrometer (Thermo Scientific). Precursor mass spectra (MS1) were acquired at 120,000 resolution from 380-1500 m/z with an automatic gain control (AGC) target of 2 x 10^5^ and a maximum injection time of 50 ms. Dynamics exclusion was set for 15 seconds with a mass tolerance of +/- 10 ppm. Quadrupole isolation for MS2 scans was 1.6 Da sequencing the most intense ions using Top Speed for a 3 second cycle time. All MS2 sequencing was performed using collision induced dissociation (CID) at 35% collision energy and scanned in the linear ion trap. An AGC target of 1 x 10^4^ and 35 second maximum injection time was used. Selected-precursor selections of MS2 scans was used to isolate the five most intense MS2 fragment ions per scan to fragment at 65% collision energy using higher energy collision dissociation (HCD) with liberated TMT reporter ions scanned in the orbitrap at 60,000 resolution (FWHM). An AGC target of 1 x 10^5^ and 240 second maximum injection time was used for all MS3 scans. All raw files were converted to mzML files and searched against the Uniprot Human database downloaded April 1, 2015 using Mascot v2.5 with cysteine carbamidomethylation as a fixed modification, methionine oxidation, and protein N-terminal acetylation were searched as variable modifications. Peptide mass tolerance was 20 ppm for MS1 and 0.5 mDa for MS2. All peptides were thresholded at a 1% false discovery rate (FDR).

### Phosphoproteomics

#### Sample preparation and phosphopeptide enrichment

SK-N-BE(2) cells were cultured in SILAC media either with ^13^C_6_^15^N_2_-lysine/^13^C_6_^15^N_4_- arginine (Lys8/Arg10) (Heavy) or Lys0 and Arg0 (Light). Two biological replicates of near confluent Heavy cells and two replicates of near confluent Light cells were treated with 20 μM CBD for 10 minutes (4 replicates), 1 hour (4 replicates) and 3 hours (4 replicates) for phosphoproteomics analysis. Cells were harvested in 4% (w/v) SDS, 100 mM Tris, pH 8.5 and boiled at 95⁰C for 5 minutes. Samples were reduced with 10 mM TCEP and alkylated with 50mM chloroacetamide, then digested using the FASP protocol, with the following modifications: an Amicon Ultra 0.5 mL 10 kD NMWL cutoff (Millipore) ultrafiltration device was used rather than a 30 kD NMWL cutoff. Tryptic peptides were cleaned a Water HLB Oasis cartridge (Waters) and eluted with 65% (v/v) ACN, 1% TFA. Glutamic acid was added to 140 mM and TiO_2_ (Titanshere, GL Sciences) was added at a ratio of 10 mg TiO_2_:1 mg tryptic peptide and incubated for 15 minutes at ambient. The phosphopeptide-bound TiO_2_ beads were washed with 65% (v/v) ACN, 0.5% TFA and again with 65% (v/v) ACN, 0.1% TFA, then transferred to a 200 μL C8 Stage Tip (Thermo Scientific). Phosphopeptides were eluted with 65% (v/v) ACN, 1% (v/v) ammonium hydroxide and lyophilized dry.

### High pH C18 fractionation of enriched phosphopeptides

Enriched phosphopeptide samples were then suspended in 20 μL 3% (v/v) acetonitrile (ACN) and 0.1% (v/v) trifluoroacetic acid (TFA) and loaded onto a custom fabricated reverse phase C18 column (0.5 x 200 mm C18, 1.8 μm 120Å Uchrom (nanoLCMS Solutions) maintained at 25⁰C and running 15 μL/min with buffer A, 10 mM ammonium formate, pH 10 and buffer B, 10 mM ammonium formate, pH 10 in 80% (v/v) ACN with a Waters M-class UPLC (Waters). Peptides were separated by gradient elution from 3% B to 50% B in 25 minutes, then from 50% B to 100% B in 5 minutes. Fractions were collected in seven rounds of concatenation for 30 sec per fraction for a final of twelve high pH C18 fractions. Samples were dried and stored at -80⁰C until analysis.

### Liquid Chromatography/Mass spectrometry analysis of phosphopeptide fractions

Samples were suspended in 3% (v/v) acetonitrile, 0.1% (v/v) trifluoroacetic acid and direct injected onto a 1.7 μm, 130 Å C18, 75 μm X 250 mm M-class column (Waters), with a Waters M-class UPLC. Tryptic peptides were gradient eluted at 300 nL/minute, from 3% acetonitrile to 20% acetonitrile in 100 minutes into an Orbitrap Fusion mass spectrometer (Thermo Scientific). Precursor mass spectrums (MS1) were acquired at 120,000 resolution from 380-1500 m/z with an AGC target of 2 x 10^5^ and a maximum injection time of 50 ms. Dynamic exclusion was set to 20 seconds with a mass tolerance of +/- 10 ppm. Isolation for MS2 scans was 1.6 Da using the quadrupole, and the most intense ions were sequenced using Top Speed for a 3 second cycle time. All MS2 sequencing was performed using higher energy collision dissociation (HCD) at 35% collision energy and scanned in the linear ion trap. An AGC target of 1 x 10^4^ and 35 second maximum injection time was used. Rawfiles were searched against the Uniprot human database using MaxQuant (version 1.6.0.13) with cysteine carbamidomethylation as a fixed modification. Methionine oxidation, protein N-terminal acetylation, and phosphorylation of serine, threonine and tyrosine were searched as variable modifications. All peptides and proteins were thresholded at a 1% false discovery rate (FDR).

### Bulk Metabolomics Sample Preparation

Cultured cells were harvested, washed with PBS, flash frozen, and stored at -80C until analysis. Prior to LC-MS analysis, samples were placed on ice and re-suspended with methanol:acetonitrile:water (5:3:2, v/v/v) at a concentration of 2 million cells per ml. Suspensions were vortexed continuously for 30 min at 4°C. Insoluble material was removed by centrifugation at 10,000 g for 10 min at 4°C and supernatants were isolated for metabolomics analysis by UHPLC-MS. This method was used for cholesterol precursors and free headgroups.

### UHPLC-MS analysis for Bulk Metabolomics

Analyses were performed as previously published (Nemkov et al., 2017, 2019). Briefly, the analytical platform employs a Vanquish UHPLC system (Thermo Fisher Scientific, San Jose, CA, USA) coupled online to a Q Exactive mass spectrometer (Thermo Fisher Scientific, San Jose, CA, USA). Samples were resolved over a Kinetex C18 column, 2.1 x 150 mm, 1.7 µm particle size (Phenomenex, Torrance, CA, USA) equipped with a guard column (SecurityGuard^TM^ Ultracartridge – UHPLC C18 for 2.1 mm ID Columns – AJO-8782 – Phenomenex, Torrance, CA, USA) (A) of water and 0.1% formic acid and a mobile phase (B) of acetonitrile and 0.1% formic acid for positive ion polarity mode, and an aqueous phase (A) of water:acetonitrile (95:5) with 1 mM ammonium acetate and a mobile phase (B) of acetonitrile:water (95:5) with 1 mM ammonium acetate for negative ion polarity mode. Samples were eluted from the column using either an isocratic elution of 5% B flowed at 250 µl/min and 25°C or a gradient from 5% to 95% B over 1 minute, followed by an isocratic hold at 95% B for 2 minutes, flowed at 400 µl/min and 30°C. The Q Exactive mass spectrometer (Thermo Fisher Scientific, San Jose, CA, USA) was operated independently in positive or negative ion mode, scanning in Full MS mode (2 μscans) from 60 to 900 m/z at 70,000 resolution, with 4 kV spray voltage, 15 sheath gas, 5 auxiliary gas. Calibration was performed prior to analysis using the Pierce^TM^ Positive and Negative Ion Calibration Solutions (Thermo Fisher Scientific). Acquired data was then converted from raw to .mzXML file format using Mass Matrix (Cleveland, OH, USA). Metabolite assignments, isotopologue distributions, and correction for expected natural abundances of deuterium, ^13^C, and ^15^N isotopes were performed using MAVEN (Princeton, NJ, USA).(Clasquin et al., 2012) Graphs, heat maps and statistical analyses (either T-Test or ANOVA), metabolic pathway analysis, PLS-DA and hierarchical clustering was performed using the MetaboAnalyst package (www.metaboanalyst.com)(Chong et al., 2018).

### Lipidomics Sample Preparation

Extraction of cholesterol, precursors, free fatty acids, cholesteryl esters, and phospholipids were performed in the following manner. SK-N-BE(2) cells in 10 cm dishes were washed with 10 mL PBS twice and then cells were scraped and pelleted at 400 rcf for 2 minutes. Cell pellets were resuspended in 100% methanol at 4°C and sonicate at 70% power in 10 pulses, 5 seconds on/5 seconds off. The resulting lysate was rotated for 60 minutes at room temperature, followed by centrifugation for 20 min at 4°C (16,100 rcf). Subcellular fractionation of organelles from intact SK-N-BE(2) cells was done in the following manner to assess subcellular CBD distribution. Cells in 10 cm culture dishes were harvest by washing twice with 10 mL PBS at room temperature, followed by trypsinization using a cell culture grade Trypsin/EDTA solution (ThermoFisher). Trypsinized cells were quenched by addition of 2 mL 10% FBS containing DMEM and cells were pelleted by centrifugation for 2 min at 4°C (200 rcf). Cell pellets were wash one time with 10 mL PBS, and resuspended in 1 mL Tween20 Buffer (1x PBS, 0.05 % Tween20, 5 mM EDTA). This lysate was subjected to mechanical disruption using a 1mL glass Dounce homogenizer, 10 full passes at 4°C. Nuclei was pelleted from homogenate by centrifugation for 5 min at 4°C (2,000 rcf). Supernatant was separated and insoluble ER membranes were pelleted by centrifugation for 10 min at 4°C (4,000 rcf). Supernatant was separated and insoluble plasma membranes were pelleted by centrifugation for 10 min at 4°C (16,000 rcf). Extraction of all fractions was done in 100% methanol for 2 hours at room temperature and rotation end-over-end, followed by removal of insoluble material by centrifugation for 20 min at 20°C (16,100 rcf).

### UHPLC-MS analysis for Lipidomics

Samples were analyzed as published (Reisz et al., 2019). Briefly, analytes were resolved over an ACQUITY HSS T3 column (2.1 x 150 mm, 1.8 µm particle size (Waters, MA, USA) using an aqueous phase (A) of 25% acetonitrile and 5 mM ammonium acetate and a mobile phase (B) of 90% isopropanol, 10% acetonitrile and 5 mM ammonium acetate. The column was equilibrated at 30% B, and upon injection of 10 μL of extract, samples were eluted from the column using the solvent gradient: 0-9 min 30-100% B and 0.325 mL/min; hold at 100% B for 3 min at 0.3 mL/min, and then decrease to 30% over 0.5 min at 0.4 ml/min, followed by a re-equilibration hold at 30% B for 2.5 minutes at 0.4 ml/min. The Q-Exactive mass spectrometer (Thermo Fisher) was operated in positive and negative ion mode using electrospray ionization, scanning in Full MS mode (2 μscans) from 150 to 1500 m/z at 70,000 resolution, with 4 kV spray voltage, 45 shealth gas, 15 auxiliary gas. When required, dd-MS2 was performed at 17,500 resolution, AGC target = 1e5, maximum IT = 50 ms, and stepped NCE of 25, 35 for positive mode, and 20, 24, and 28 for negative mode. Calibration was performed prior to analysis using the PierceTM Positive and Negative Ion Calibration Solutions (Thermo Fisher). Acquired data was then converted from .raw to .mzXML file format using Mass Matrix (Cleveland, OH, USA). Samples were analyzed in randomized order with a technical mixture injected incrementally to qualify instrument performance. This technical mixture was also injected three times per polarity mode and analyzed with the parameters above, except CID fragmentation was included for unknown compound identification. Metabolite assignments were made based on accurate intact mass (sub 5 ppm), isotope distributions, and relative retention times, and comparison to analytical standards in the SPLASH Lipidomix Mass Spec Standard (Avanti Polar Lipids) using MAVEN (Princeton, NJ, USA). Discovery mode analysis was performed with standard workflows using Compound Discoverer and Lipid Search 4.0 (Thermo Fisher Scientific, San Jose, CA).

### Confocal Microscopy of Cholesterol and Lysosomes

SK-N-BE(2) cells were seeded into fibronectin coated glass bottom 96 well plates (Matriplate) at a cell density of 40,000 cells/well using low background imaging media (FluoroBrite DMEM with all supplements described, above). At the time of seeding, lysotracker Deep Red (ThermoFisher) was added at a 1000x dilution and NBD-cholesterol (ThermoFisher) was added at a final concentration of 10 µg/ mL. After 24 hrs, CBD or ethanol vehicle was added to a final concentration of 20 µM and incubated for an additional 24 hrs prior to imaging using a Nikon A1R laser scanning confocal microscope for acquisition with the FITC and TRITC channels. In experiments using U18666A, a final concentration of 10 µg/ml was used and was added simultaneously with CBD.

### Assaying Cell Viability and Apoptosis

Cell viability for SK-N-BE(2) cells was conducted using a fluorometric cell viability assay using Resazurin (PromoKine) according to the manufacturer’s instructions.

Measurement of percent apoptotic cells was done in 384 well imaging plates (Corning #3985) seeded with 2,000 cells/well and stained with Hoescht 33258 (1µg/mL) and CellEvent Caspase-3/7 Green Detection Reagent (ThermoFisher) at a dilution of 1000x. Dyes were added at the time of seeding, 18-24 hours prior to performing experiments. For experiments using atorvastatin, atorvastatin was added 24 hrs prior to addition of CBD. For experiments involving 25-hydroxy cholesterol, U18666A, and VULM 1457, inhibitors were added simultaneously with CBD. Experiments were performed using an ImageXpress MicroXL microscope and a 10x objective, where images were acquired for each well at the indicated time points using DAPI and FITC filter sets. Using MATLAB, images were processed with custom written scripts (available upon request) that perform flatfield & background correction, identification of all cells (DAPI channel) using standard threshold above background techniques, and identification of apoptotic cells using a similar method in the FITC channel. Percent apoptotic cells was calculated from the sum of apoptotic cell pixels divided by the sum of all cell pixels for each field of view. Error displayed is the standard deviation from between 2 and 4 biological replicates.

### Fluorometric Cholesterol Oxidase Experiments and SUV preparation

SUVs were prepared by dissolving 10 mg L-α-Phosphatidylcholine (Sigma P3556) in 100 µL chloroform in a glass vial, followed by removal of solvent under vacuum at room temperature for 1 hour. For experiments using cholesterol containing SUVs, 0.74 mg of cholesterol (Sigma C8667) was mixed with 10 mg L-α-Phosphatidylcholine prior to removal of chloroform solvent. Following solvent removal, 100 µL PBS was added and a microtip sonicator was inserted to perform sonication at 70% power, 10 pulses, 5 seconds on/off at room temperature. SUVs in suspension were brought to a volume of 1 mL with addition of PBS. The resulting SUVs in suspension were used at a dilution of 100-fold in subsequent cholesterol oxidase reactions. Cholesterol oxidase reactions were performed using reagents from the Amplex Red Cholesterol Assay Kit (ThermoFisher A12216), where each reaction was performed in 50 µL volumes using: 0.5 µL SUVs solution, 0.05 µL cholesterol oxidase solution, 0.05 µL HRP solution, 0.05 µL Amplex Red/DMSO made according to manufacturer’s instructions, and the indicated CBD concentrations in PBS. Reaction volume was brought to 50 µL using PBS. In cases where SUVs were not used, either 1 µg/reaction 25-OH cholesterol or 5 µg/reaction MBCD:Cholesterol (1:2) was substituted for SUVs. MBCD:Cholesterol was prepared as previously described (Widenmaier et al., 2017). Cholesterol oxidase reactions were performed in Corning 384 well optical imaging plates (#3985) in an ImageXpress MicroXL widefield fluorescence microscope using the TRITC filter sets, where 1 ms exposure time images were taken of each well 20 µm above the well bottom every 10 minutes for 5 hours at 37 °C, using a 10x objective. Images were flatfield corrected and the sum of fluorescence intensity across all 540x540 pixels was calculated using custom MATLAB scripts that are available upon request. Product formation of Amplex red was found to be linear within 0-1 hours, and data between t=0 and t=1 hours was used to calculate the average rate of increase in TRITC fluorescence using Microsoft Excel. Displayed error bars represent the standard deviation of three or more replicate reactions.

### FRAP Experiments Using Synthetic Membranes

The formulation and techniques to create SUVs were repeated with addition of 108 µg of 22-NBD cholesterol to L-α-Phosphatidylcholine and cholesterol prior to the solvent removal step described for preparation of SUVs. A 1:5 dilution of NBD-cholesterol containing SUV suspension:PBS was added to each well of glass bottom 96 well plates (MatriPlate MGB096-1-2-LG-L) such that each well contains 100 µl of diluted SUV suspension. 96 well plates were exposed to centrifugation for 20 minutes at 2000 rcf using a swinging bucket rotor at room temperature. A microtip sonicator was inserted into each well to perform sonication at 20% power, 20 pulses, 2 seconds on/off at room temperature. The contents of each well were washed three times with 150 µl of PBS, and subsequent experiments were performed with 250 µl PBS containing ethanol vehicle, 20 µM CBD and/or 20 µM DHA. CBD and DHA were incubated in wells for 1 hr at room temperature prior to imaging and FRAP experiments using a Nikon A1R microscope. Photobleaching was performed using Nikon Elements software with the following parameters: framerate 250 ms, 100% power 488 laser for photobleaching for 250 ms, and optical settings for FITC. Analysis was performed using ImageJ and Microsoft Excel. All trends were normalized by division of mean intensity within the photobleached region to a region of identical size remote from the photobleached region. Error bars indicate the standard error of the mean from three replicates.

### Seahorse Extracellular Flux Analysis

Oxygen consumption rate and extracellular acidification rate were measured using the SeahorseXF^e^24 Extracellular Flux Analyzer and the Agilent Seahorse XF Cell Energy Phenotype Test Kit. Cells were plated at 2×10^4 cells per well in XF^e^24 microplates. Cells were treated with either 20 µM CBD or ethanol as a vehicle control either 24 hours or 2 hours prior to assaying. The day of the assay cells were washed with an assay medium containing 20 µM CBD or vehicle and placed at 37°C in a CO2 free incubator for 1 hour. 1 µM Oligomycin and 1 µM FCCP were injected by the Seahorse analyzer as oxygen consumption rate and extracellular acidification rate were measured per manufacturer’s protocol.

### Methods for Supplemental Figure S6

40,000 SK-N-BE(2) cells were seeded into each well of a 96 well plate. After 18 hours, cells were exposed to vehicle or CBD for 24 hours, followed by 1-hour incubation with Filipin (Sigma F9765) in the presence of Hoescht33258 and Propidium Iodide. Cells were imaged using DAPI and TRITC filter sets in an ImageXpress MicroXL microscope. TRITC Fluorescence was quantified as the sum of pixel intensity above background after flatfield correction.

## Supporting information

Metabolomics time course data

Phosphoproteomics differential statistical analysis of phospho-sites

Subcellular proteomics, cytoplasmic fraction, differential statistical analysis of proteins

Subcellular proteomics, nuclear fraction, differential statistical analysis of proteins

Subcellular proteomics, PM fraction, differential statistical analysis of proteins

RNA-seq differential statistical analysis of mRNA transcripts

Lipidomics data

## Abbreviations

25-OHC: 25-hydroxycholesterol
ACACA: acetyl-CoA carboxylase
ACAT: acyl-coenzyme A cholesterol O-acyltransferase
AMPK: AMP-activated protein kinase
AMKAR: AMP-activated protein kinase activity reporter
APOB: apolipoprotein B
APOE: apolipoprotein E
ATF4: activating transcription factor 4
CBD: cannabidiol
CAMKKß: Ca^2+^/ calmodulin-dependent protein kinase kinase ß
CaV3: voltage gated calcium channel 3
CREB: cAMP response element-binding protein
DMSO: dimethylsulfoxide
DHA: docosahexaenoic acid
DHCR24: 24-dehydrocholesterol reductase
EEF2: eukaryotic elongation factor 2
EEF2K: eukaryotic elongation factor 2 kinase
ERK: extracellular signal-regulated kinase
ECAR: extracellular acidification rate
ER: endoplasmic reticulum
FCCP: trifluoromethoxy carbonylcyanide phenylhydrazone
FDA: Food and Drug Administration
FRAP: fluorescence recovery after photobleaching
FRET: Förster resonance energy transfer
GPR55: G protein-coupled receptor 55
GRAMD1: GRAM domain containing protein 1
GTP: guanosine triphosphate
HK1: Hexokinase 1
HMGCR: 3-hydroxy-3-methylglutaryl-CoA reductase
HSP60: heat shock protein 60
LDLR: low density lipoprotein receptor
LPE: lysophosphatidylethanolamine
MBCD: methyl ß cyclodextrin
MCMC: Markov chain Monte Carlo
NaV1.1: sodium channel, voltage-gated, type I, alpha subunit
NFE2L2: nuclear factor erythroid 2-related factor 2
NPC1: Niemann-Pick disease, type C1
OCR: oxygen consumption rate
PCA: principal component analysis
PC: phosphatidylcholine
PE: phosphatidylethanolamine
SCAP: SREBP cleavage-activating protein
SGPL1: Sphingosine-1-phosphate lyase
SILAC: stable isotope labeling of amino acids in culture
SOD1: superoxide dismutase 1
SP1: specificity protein 1 transcription factor
SREBP: sterol regulatory element-binding protein
STK11: serine/ threonine kinase 11
SUV: synthetic unilamellar vesicles TCA tricarboxylic acid cycle
TMT: tandem mass tag
TRPM8: transient receptor potential cation channel subfamily M member 8
TRPV: transient receptor potential
VDAC1: voltage-dependent anion channel 1

## Acknowledgements

The authors acknowledge the BioFrontiers Computing Core at the University of Colorado Boulder for providing High Performance Computing resources (NIH 1S10OD012300) supported by BioFrontiers IT. The imaging work was performed at the BioFrontiers Institute Advanced Light Microscopy Core. The Molecular Devices ImageXpress was supported by NIH grant 1S10RR026680-01A1. Laser scanning confocal microscopy was supported by NIST-CU Cooperative Agreement award number 70NANB15H226. This work was supported by a DARPA cooperative agreement, 13-34-RTA-FP-007, to WMO, MHBS, XL, and AD.

## Authorship Contributions

**Steven E. Guard:** Formal Analysis, Investigation, Visualization, Writing – Original Draft, Writing – Review & Editing; **Douglas A. Chapnick:** Conceptualization, Investigation, Methodology, Visualization, Writing – Original Draft; **Zachary C. Poss:** Investigation; **Christopher C. Ebmeier:** Investigation, Visualization; **Jeremy Jacobsen:** Formal Analysis, Methodology, Visualization; **Travis Nemkov:** Investigation, Visualization; **Kerri A. Ball:** Investigation; **Kristofor J. Webb:** Investigation; **Helen L. Simpson:** Investigation; **Stephen Coleman:** Formal Analysis, Methodology, Visualization; **Eric Bunker:** Visualization; **Adrian Ramirez:** Investigation; **Julie A. Reisz:** Investigation**; Robert Sievers:** Resources**; Michael H.B. Stowell:** Funding Acquisition, Supervision; **Angelo D’Alessandro:** Acquisition of Funding, Supervision; **Xuedong Liu:** Conceptualization, Investigation, Methodology, Funding Acquisition, Supervision; **William M. Old:** Conceptualization, Funding Acquisition, Investigation, Methodology, Project Administration, Writing-Original Draft, Writing – Review & Editing

## Resource Availability

All unique/stable reagents generated in this study are available from the lead contact, William Old with a completed Materials Transfer Agreement.

## Data and Code Availability Statements

Proteomics and phosphoproteomics raw data is available at the MassIVE repository ID MSV000085479 accessible at https://doi.org/doi:10.25345/C5571V

Source data for RNA-seq experiments is accessible at GEO with the identifier GSE151512 at https://www.ncbi.nlm.nih.gov/geo/query/acc.cgi?acc=GSE151512

The code generated during this study is available at GitHub, DOI:10.5281/zenodo.3861043, URL: https://github.com/CUOldLab/cbd-manuscript-code-guardse-2020

## Supplemental Tables Summary

**Table S1.** Global Metabolomics Results, Related to Figure 1C and Supplementary Figures Throughout

**Table S2.** Phosphoproteome Results, Related to Figure 2

**Table S3.** Cytosolic Proteome Results, Related to Figure 3

**Table S4.** Nuclear Proteome Results, Related to Figure 3

**Table S5.** Membrane Proteome Results, Related to Figure 3

**Table S6.** RNA-seq Differential Analysis Results, Related to Figure 3D

**Table S7.** Lipidomics Results, Related to Figure 4

