## Supplemental Table 1 and Supplemental Figures 1-6 for "Multi-Omic Analysis Reveals Disruption of Cholesterol Homeostasis by Cannabidiol in Human Cell Lines"

**Supplemental Information Legends:**

**Supplemental Table 1. Biosensors used for profiling the EC50 Effects of CBD.**

The name of each genetically encoded biosensor gene used to profile diverse activities in cell lines is displayed along with its target analyte and literature source.

**Supplemental Figure 1. FRET-based biosensor screening reveals dose dependent molecular responses to CBD**

**(A)** Genetically encoded FRET biosensors for diverse biochemical pathways were delivered to both SK-N-BE(2) and HaCaT Cells. Heatmaps display normalized FRET ratio over 20 minute time intervals from 0.343 μM to 100 μM CBD treatment.

**Supplemental Figure 2. CBD-dependent phosphorylation changes are enriched in AMPK substrate motifs**

**(A)** Volcano plot of phosphorylation sites identified by LC/MS 10 minutes post treatment with 20 μM CBD. 5 proteins were identified with adjusted $p < 0.05$ and $FC > |0.5|$ by 10 minutes (text in white box / red dot label) **(B)** BioPlanet Pathway enrichment of significantly changing phosphorylation sites at 1 and 3 hours post CBD treatment. * adjusted $p < 0.05$; ** adjusted $p < 0.01$ **(C-D)** Motif enrichment tables for significantly changing phosphorylated sites against global background at 1 and 3 hours post treatment with 20 μM CBD. **(E)** Abundance of *de novo* synthesized and elongated fatty acids are shown for $^{13}C_2$-butanoic acid (C4) and $^{13}C_2$-octadecenoic acid (C18:1) in both vehicle-treated (blue) and CBD-treated (20 μM) cells (red) Lines represent median values and shaded areas covered standard error of the median **(F)** The relative abundance of acylcarnitines (AC) are shown as a heat map with abundance plotted by Z-score according to a color gradient from blue to red for lowest to highest abundance. The acyl-chain length is described in parentheses. **(G)** The relative levels of $^{13}C_2$ incorporation into lactate and TCA cycle intermediates in both vehicle-treated (blue) and CBD-treated cells (red).

**Supplemental Figure 3. pH-dependent subcellular fractionation generates compositionally distinct proteome fractions**

**(A)** Principal component analysis (PCA) of proteome fractions in a CBD time course based on bioconductor EDGE analysis of 5 replicate pairs as described in the STAR METHODS **(B)** PCA of proteome fractions and whole cell lysate in untreated conditions. Each dot represents a biological replicate. **(C-E)** Cellular component enrichment analysis of proteins identified in each of the proteome fractions. Significantly enriched gene ontologies from each fraction are plotted by log of the false discovery rate (FDR) versus enrichment ratio. Dot color is scaled on the number of proteins found within each enrichment. **(F)** Differential expression for HK1 between membrane and nuclear fractions over time. **(G)** Relative abundance of Hexose phosphate (representing a pool of sugar phosphates largely including glucose 6-phosphate) and reduced glutathione is shown. Lines represent median values and shaded areas covered standard error of the median. **(H)** Upstream regulator analysis of differentially expressed transcripts measured by RNAseq in CBD treated cells. Number of genes enriched at each timepoint are displayed on the right.

**Supplemental Figure 4. CBD treatment results in increased flux of biosynthetic precursors by 24 hours**

**(A-B)** Plots of $^{13}C$ labeled and total + labeled abundances of cholesterol biosynthesis metabolites measured in methanol extracts from SK-N-BE(2) cells treated with vehicle or 20 μM CBD. **(C)** Fluorometric measurement of total cellular cholesterol from methanol extracts of SK-N-BE(2) cells using the Amplex Red Cholesterol Assay Kit. **(D)** Ethanolamine phosphate abundance time course from global metabolomics profiling experiment.

**Supplemental Figure 5. CBD-dependent apoptosis is cholesterol dependent across cell lines**

**(A)** HACAT cells were assessed for apoptosis at 24 hours using live cell microscopy using a resazurin based fluorometric cell viability stain. Cells were treated with 10 μM atorvastatin and exposed to increasing doses of CBD. **(B)** HACAT cells were treated with 20 μM CBD and exposure to increasing doses of 25-OH cholesterol. **(C)** SK-N-BE(2) and HEK293T cells were treated with different combinations of 20 μM CBD, 10 μM U18666A, 5μM VULM and 15μg/ ml 25- OH Cholesterol and were measured for apoptosis rates over time.

**Supplemental Figure 6. CBD alters cholesterol orientation in complex membranes**

**(A)** Synthetic small unilamellar vesicles (SUVs) free of cholesterol and composed of 100% Phosphatidylcholine were analyzed with a fluorogenic cholesterol oxidase reaction to determine the effect of CBD on initial reaction rate in the presence and absence of cholesterol oxidase enzyme. **(B)** 1μg/ reaction of free 25-OH cholesterol was used in the absence of SUVs in a cholesterol oxidase fluorogenic assay to measure reaction rate with non-membrane resident cholesterol. **(C)** ER derived vesicles were used as a source of cholesterol for similar experiments as displayed in suppl 6B. **(D-E)** SK-N-BE(2) and HEK293T cells were treated with increasing amounts of DHA and evaluated for apoptosis. DHA (75 μM) was then evaluated for rescue by either low dose CBD (6.25 μM) or MBCD (300 nM). **(F)** Fluorescent measurement of propidium iodide uptake in live cells treated with 20 μM CBD and a dose curve of Filipin at 24 hours. CBD increased membrane permeability to propidium iodide.

| Targeted Analyte | Biosensor Gene Name | Original Literature Source |
|---|---|---|
| Adam17 Protease Activity | TSEN | Chapnick DA, Bunker E, Liu X (2015) A biosensor for the activity of the "sheddase" TACE (ADAM17) reveals novel and cell type-specific mechanisms of TACE activation. Science signaling 8 (365) |
| AMP Kinase Activity | AMPKAR | Tsou P, Zheng B, Hsu CH, Sasaki AT, Cantley LC (2011) A fluorescent reporter of AMPK activity and cellular energy stress. Cell metabolism 13 (4):476-486 |
| ATP Abundance | ATEAM | Imamura H, Nhat KP, Togawa H, Saito K, Iino R, Kato-Yamada Y, Nagai T, Noji H (2009) Visualization of ATP levels inside single living cells with fluorescence resonance energy transfer-based genetically encoded indicators. Proceedings of the National Academy of Sciences of the United States of America 106 (37):15651-15656 |
| Cytosolic $Ca^{2+}$ Abundance | D3-cpv | Ravier MA, Cheng-Xue R, Palmer AE, Henquin JC, Gilon P (2010) Subplasmalemmal Ca(2+) measurements in mouse pancreatic beta cells support the existence of an amplifying effect of glucose on insulin secretion. Diabetologia 53 (9):1947-1957 |
| Cytosolic ERK Activity | EKAR-NES | Komatsu N, Aoki K, Yamada M, Yukinaga H, Fujita Y, Kamioka Y, Matsuda M (2011) Development of an optimized backbone of FRET biosensors for kinases and GTPases. Molecular biology of the cell 22 (23):4647-4656 |
| ER $Ca^{2+}$ Abundance | D1ER | Palmer AE, Jin C, Reed JC, Tsien RY (2004) Bcl-2-mediated alterations in endoplasmic reticulum Ca2+ analyzed with an improved genetically encoded fluorescent sensor. Proceedings of the National Academy of Sciences of the United States of America 101 (50):17404-17409 |
| Glucose Abundance | FLIPglu-30uDelta13V | Takanaga H, Frommer WB (2010) Facilitative plasma membrane transporters function during ER transit. FASEB journal : official publication of the Federation of American Societies for Experimental Biology 24 (8):2849-2858 |

| | | |
|---|---|---|
| Glutamine Abundance | FLIPQTV3.0 8m | Gruenwald K, Holland JT, Stromberg V, Ahmad A, Watcharakichkorn D, Okumoto S (2012) Visualization of glutamine transporter activities in living cells using genetically encoded glutamine sensors. PloS one 7 (6) |
| Lactate Abundance | Laconic | San Martin A, Ceballo S, Ruminot I, Lerchundi R, Frommer WB, Barros LF (2013) A genetically encoded FRET lactate sensor and its use to detect the Warburg effect in single cancer cells. PloS one 8 (2) |
| mTOR Kinase Activity | TORCAR | Zhou X, Clister TL, Lowry PR, Seldin MM, Wong GW, Zhang J (2015) Dynamic Visualization of mTORC1 Activity in Living Cells. Cell reports |
| PKD Kinase Activity | DKAR | Kunkel MT, Toker A, Tsien RY, Newton AC (2007) Calcium-dependent regulation of protein kinase D revealed by a genetically encoded kinase activity reporter. The Journal of biological chemistry 282 (9):6733-6742 |
| Plasma Membrane Electrostatic Potential (Charge) | MCS+ | Ma Y, Yamamoto Y, Nicovich PR, Goyette J, Rossy J, Gooding JJ, Gaus K (2017) A FRET sensor enables quantitative measurements of membrane charges in live cells. Nature biotechnology 35 (4):363-370 |
| Plasma Membrane Potential | VSFP-CR | Lam AJ, St-Pierre F, Gong Y, Marshall JD, Cranfill PJ, Baird MA, McKeown MR, Wiedenmann J, Davidson MW, Schnitzer MJ, Tsien RY, Lin MZ (2012) Improving FRET dynamic range with bright green and red fluorescent proteins. Nature methods 9 (10):1005-1012 |
| Pyruvate Abundance | Pyronic | San Martin A, Ceballo S, Baeza-Lehnert F, Lerchundi R, Valdebenito R, Contreras-Baeza Y, Alegria K, Barros LF (2014) Imaging mitochondrial flux in single cells with a FRET sensor for pyruvate. PloS one 9 (1) |

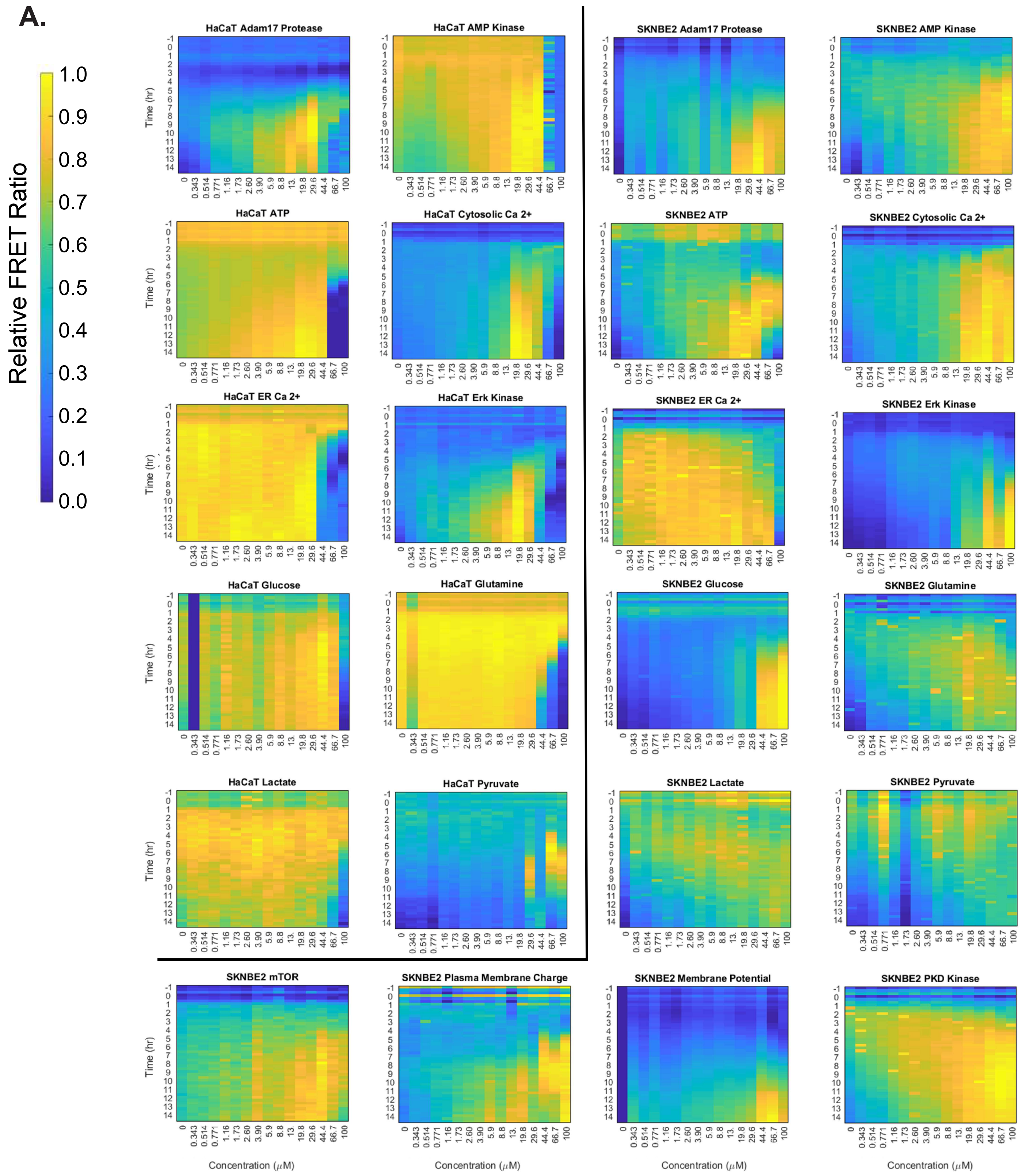


### Supplemental Figure 1

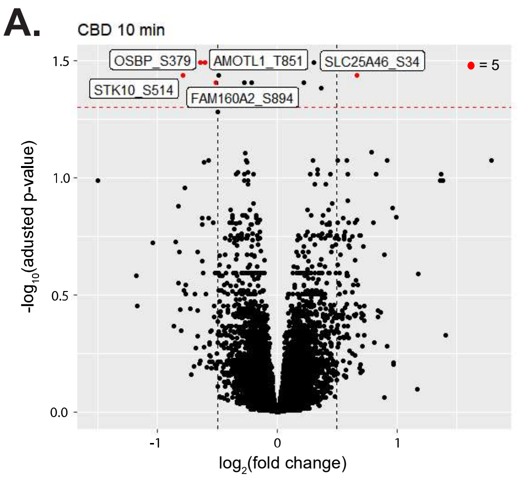


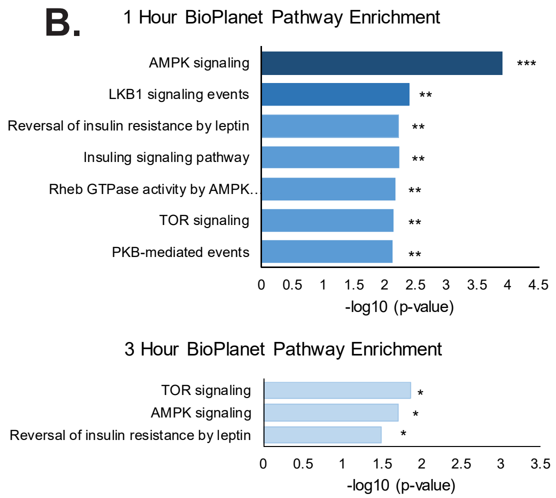


**C.** **1 Hour post-CBD Treat**

| Id | Motif | ZScore | P-Value | Fold Change | Foreground Matches | Foreground Size | Background Matches | Background Size |
|---|---|---|---|---|---|---|---|---|
| 1 | .G..St.S.R. | 10.43 | 0.00E+00 | 3366.33 | 2 | 20 | 15 | 504949 |
| 2 | **..R..s.....** | 8.98 | 0.00E+00 | 5.26 | 30 | 105 | 43563 | 802214 |
| 3 | **..R..s...L.** | 7.74 | 5.11E-15 | 16.03 | 8 | 105 | 3813 | 802214 |
| 4 | .GGS.t..... | 7.55 | 2.11E-14 | 257.63 | 2 | 20 | 196 | 504949 |
| 5 | .GG..t.L... | 7.51 | 2.99E-14 | 248.74 | 2 | 20 | 203 | 504949 |
| 6 | P..S.t..P.. | 6.93 | 2.13E-12 | 159.79 | 2 | 20 | 316 | 504949 |
| 7 | ...S.t..P.. | 6.58 | 2.35E-11 | 31.93 | 4 | 20 | 3163 | 504949 |
| 8 | ..R..s..P.. | 6.55 | 2.96E-11 | 14.87 | 6 | 105 | 3082 | 802214 |
| 9 | **L.R..s.....** | 6.49 | 4.15E-11 | 11.94 | 7 | 105 | 4481 | 802214 |
| 10 | ...S.t.SP.. | 6.47 | 4.93E-11 | 112.96 | 2 | 20 | 447 | 504949 |

**D.** **3 Hour post-CBD Treat**

| Id | Motif | ZScore | P-Value | Fold Change | Foreground Matches | Foreground Size | Background Matches | Background Size |
|---|---|---|---|---|---|---|---|---|
| 1 | **..R..s.....** | 9.03 | 0.00E+00 | 5.31 | 30 | 104 | 43563 | 802214 |
| 2 | **..R..s....L** | 6.75 | 7.54E-12 | 13.17 | 7 | 104 | 4099 | 802214 |
| 3 | ..R..s..S.. | 6.34 | 1.18E-10 | 11.22 | 7 | 104 | 4813 | 802214 |
| 4 | ...S.s...L. | 5.35 | 4.51E-08 | 7.63 | 7 | 104 | 7076 | 802214 |
| 5 | R....t..F.. | 5.15 | 1.33E-07 | 40.61 | 2 | 30 | 829 | 504949 |
| 6 | .....t.NK.. | 4.94 | 3.91E-07 | 34.88 | 2 | 30 | 965 | 504949 |
| 7 | **L....s.....** | 4.86 | 5.88E-07 | 2.56 | 25 | 104 | 75396 | 802214 |
| 8 | L.R..t..... | 4.82 | 7.25E-07 | 17.01 | 3 | 30 | 2968 | 504949 |
| 9 | ....Nt.D... | 4.79 | 8.28E-07 | 31.29 | 2 | 30 | 1076 | 504949 |

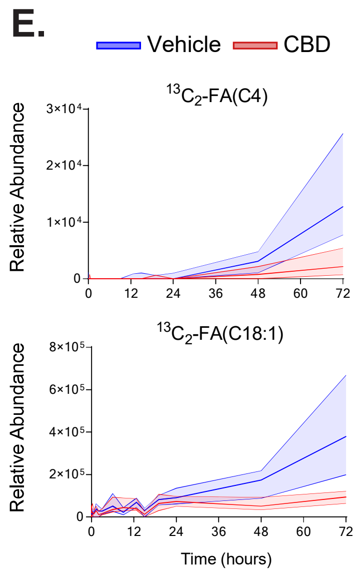


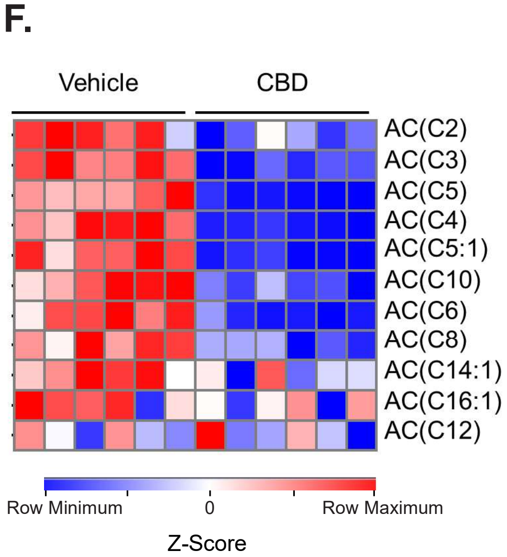


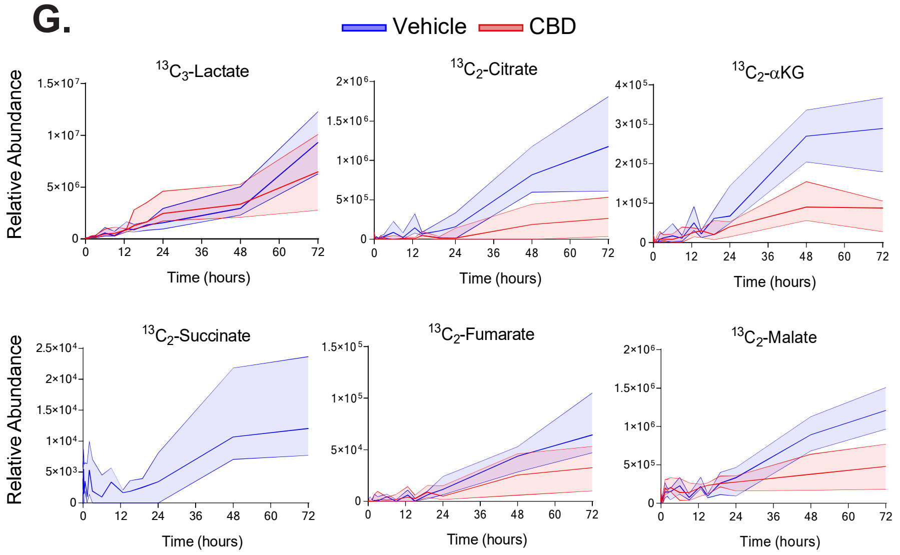


### Supplemental Figure 2

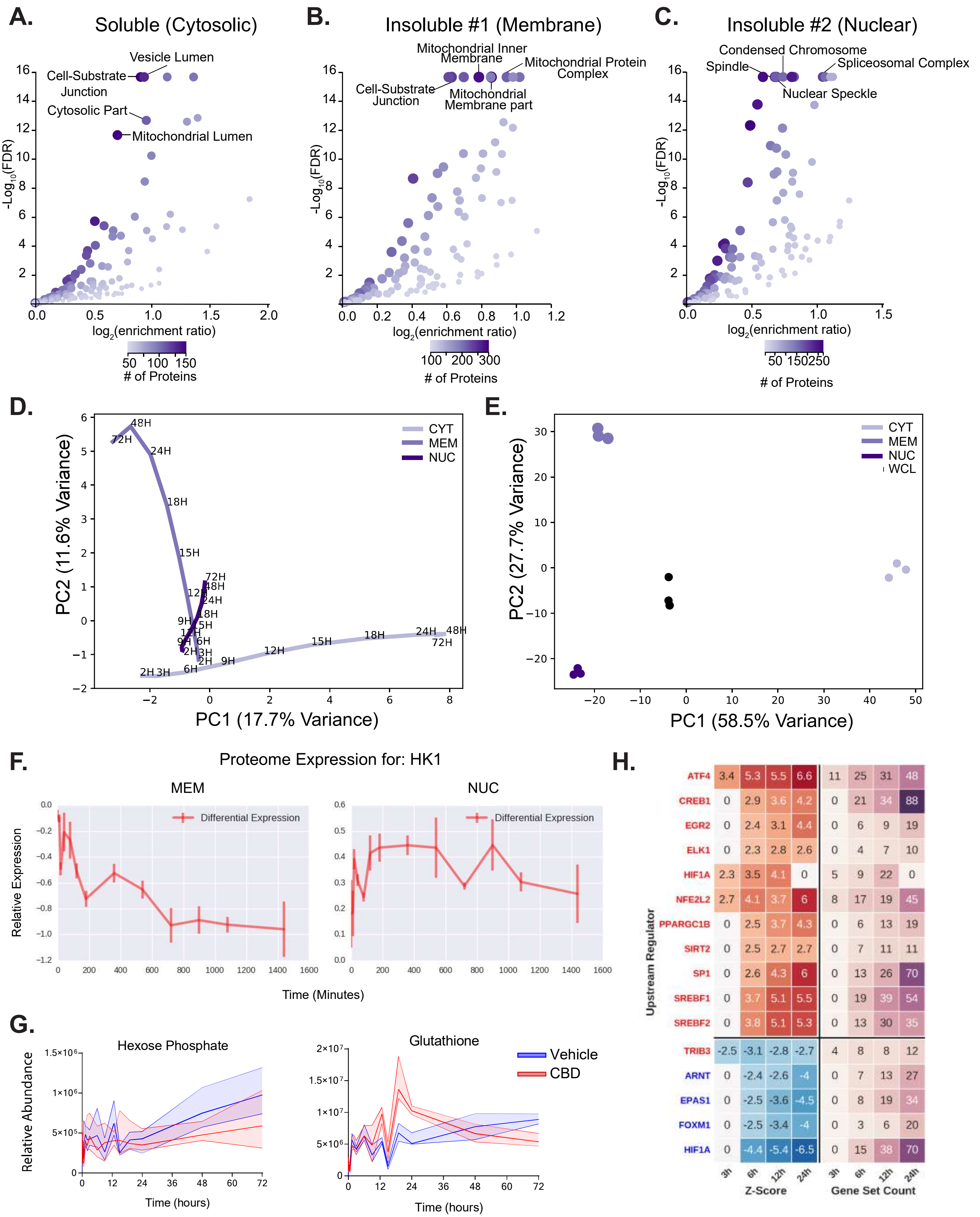


### Supplemental Figure 3

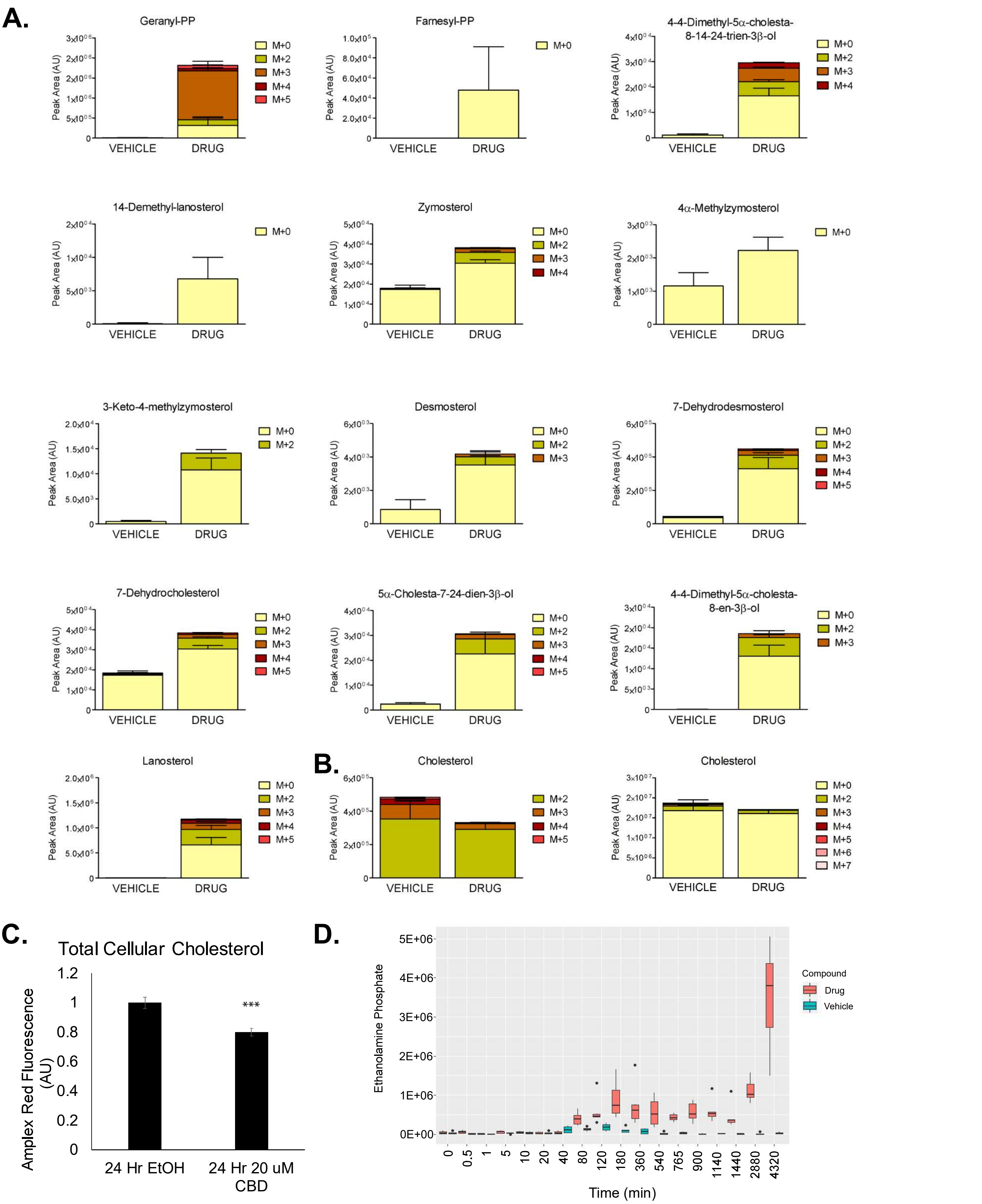


Supplemental Figure 4

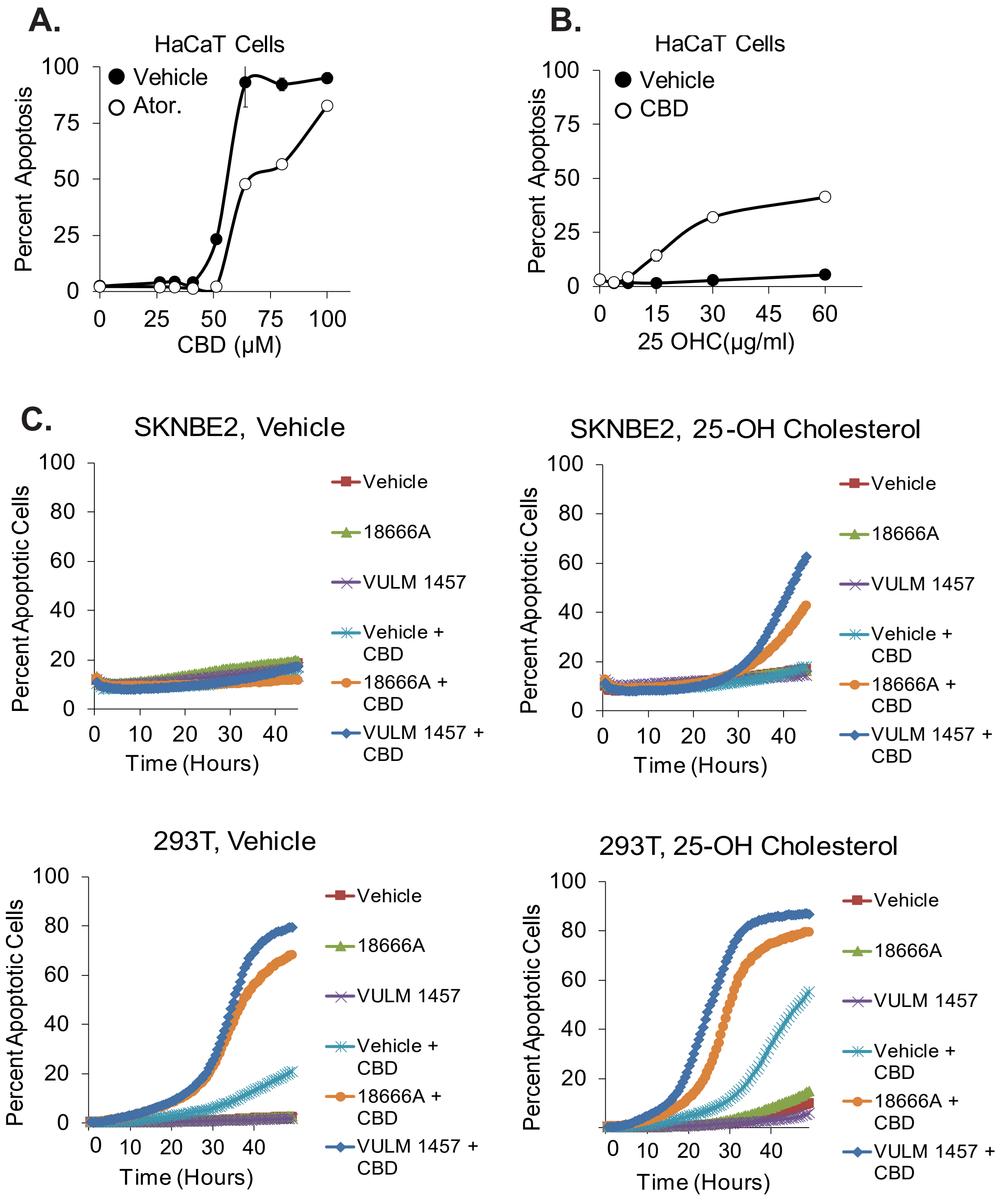


Supplemental Figure 5

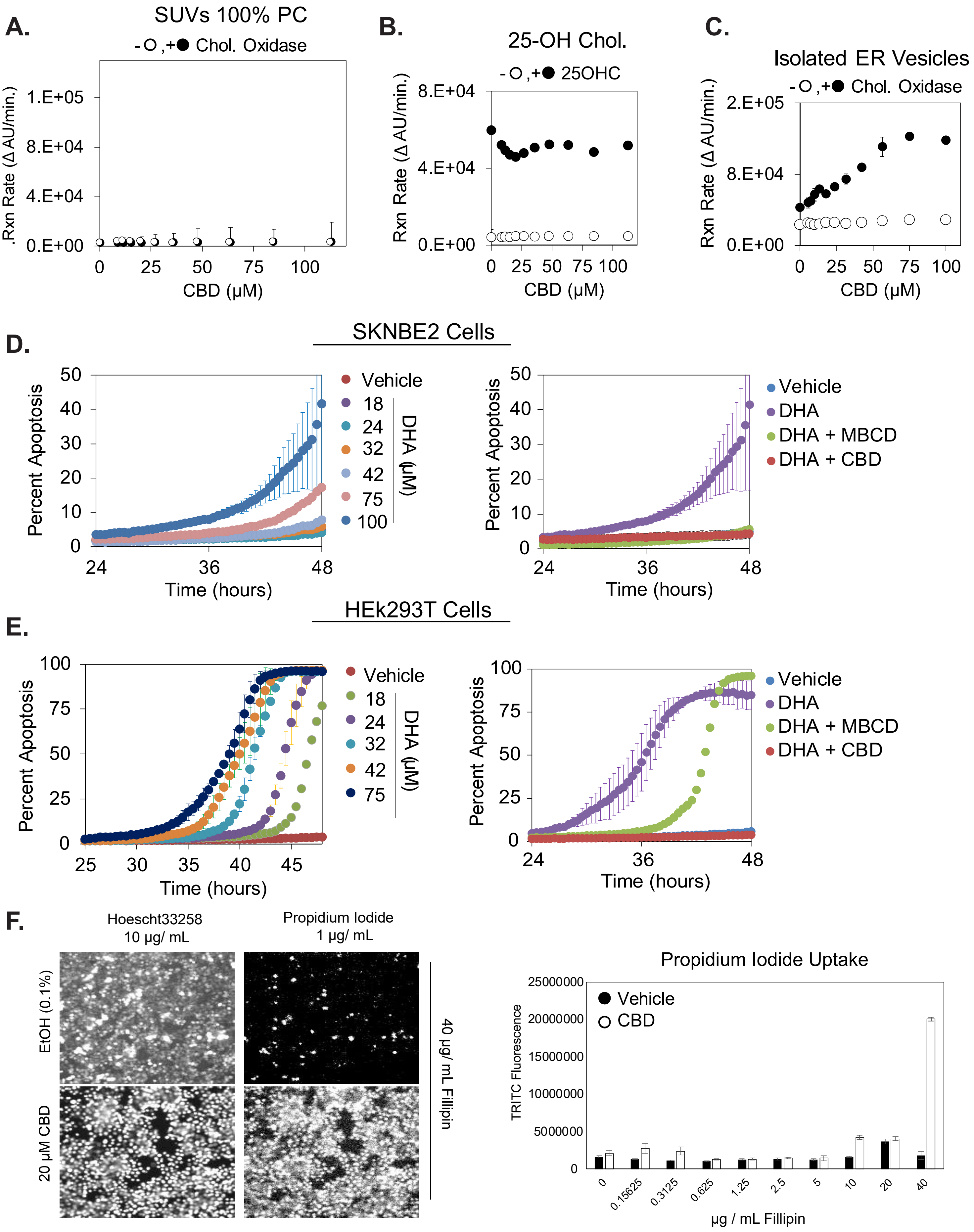


### Supplemental Figure 6
